# Disrupting the CmP signaling network unveils novel biomarkers for triple negative breast cancer in Caucasian American women

**DOI:** 10.1101/2021.09.13.460145

**Authors:** Johnathan Abou-Fadel, Muaz Bhalli, Brian Grajeda, Jun Zhang

## Abstract

Triple-negative breast cancer (TNBC) constitutes ∼15 percent of all diagnosed invasive breast cancer cases with limited options for treatment since immunotherapies that target the ER, PR and HER2 receptors are ineffective. Progesterone (PRG) is capable of inducing its effects through either classic, non-classic, or combined responses by binding to classic nuclear PRG receptors (nPRs) or non-classic membrane PRG receptors (mPRs). Under PRG-induced actions, we previously demonstrated that the CSC (CCM signaling complex) can couple both nPRs and mPRs into a CmPn signaling network which plays an important role in nPR(+) breast cancer tumorigeneses. We recently defined the novel CmP signaling network in TNBC cells, which overlapped with our previously defined CmPn network in nPR(+) breast cancer cells. In this study, we were able to demonstrate alterations to key tumorigenesis pathways in Caucasian American Women (CAW)-TNBC cells, under mPRs-specific steroid actions. These results suggest that even though TNBC diagnoses in AAW are associated with more aggressive forms of the disease, and experience a higher mortality rate, TNBC in CAW share similar altered signaling pathways, under mPRs-specific steroid actions, demonstrating the overall aggressive nature of TNBCs, regardless of racial differences. Furthermore, in this report, we have identified 21 new CAW-TNBC specific candidate biomarkers that reinforce the definitive role of the CmP signaling network in TNBC tumorigenesis, initially identified in our previous studies with AAW-TNBCs. This new set of potential prognostic biomarkers may revolutionize molecular mechanisms and currently known concepts of tumorigenesis in CAW-TNBCs, leading to hopeful new therapeutic strategies.

## Introduction

Breast cancer is the most common cancer among women in the United States and remains the second leading cause of cancer death in the world [1, 2]. Triple-negative breast cancer (TNBC) constitutes only ∼15 percent of all diagnosed invasive breast cancer cases in the United States [3, 4]. TNBC refers to breast cancer cells that lack the expression of the estrogen receptor (ER), nuclear progesterone receptor (nPR), and human epidermal growth factor receptor 2 (HER2) [5–7]. TNBCs tend to behave more aggressively and have a poorer prognosis than other subtypes of breast cancer [8–10, 11 ]. This results, in part, due to limited treatment options available compared to other molecular subtypes [12, 13]. Although not synonymous, TNBC clinical phenotypes are mostly made up of basal-like molecular subtypes and the most common histological feature is infiltrating ductal carcinoma [14]. TNBCs can exhibit geographic necrosis, a pushing border of invasion, and a stromal lymphocytic response [15]. In mammograms, TNBCs, lack speculated margins, irregular shape, and suspicious calcifications associated with other forms of breast cancers, despite being significantly larger than other subtypes at the time of diagnosis [16]. Although ultrasound and MRI are highly sensitive in detecting breast cancers, both are associated with a high false-positive rate [17]. Unfortunately, imaging studies do not provide early detection of TNBCs. Most TNBCs are detected after palpating a large mass in the breast tissue, or when the patient starts showing symptoms like tenderness of the breast and pathologic nipple discharge [18]. After a diagnosis of TNBC, patients have limited options for treatment since immunotherapies that target the ER, PR and HER2 receptors are ineffective [19]. Two of the only main options remaining are surgery and chemotherapy. If TNBCs are caught during the early-stage (stage I-III), and the tumor is small enough, it can be removed by surgery followed by an examination of the sentinel lymph node [20]. If the sentinel lymph node is found to have cancer, surgery is frequently complemented with radiation. For late-stage (stage IV) TNBCs, chemotherapy with drugs like anthracyclines, taxanes and platinum are usually first-line treatment options [21–23]. If the TNBC tumor expresses PDL1 (used to suppress immune responses), immunotherapy plus chemo are frequently implemented [24].

Progesterone (PRG), a sex steroid hormone, is capable of inducing its effects through either classic, non-classic, or combined responses by binding to either classic nuclear PRG receptors (nPRs) or non-classic membrane PRG receptors (mPRs). Under PRG-induced actions, it has been found that the CSC (CCM signaling complex), composed of CCM1, CCM2, and CCM3, can couple both nPRs and mPRs into a CSC-mPRs-PRG-nPRs (CmPn) signaling network which plays an important role in breast cancer tumorigeneses [25]. Mifepristone (MIF), an nPR antagonist and mPR agonist, has been shown to suppress basal TNBC stem cells [26]. Our previous data show that combined actions (MIF+PRG) is mPR specific, and work synergistically towards mPRs [25]. Furthermore, several reports have demonstrated that elevated levels of MIF can enhance growth inhibition and induction of apoptosis in the presence of high dose progesterone in either nPR(+/-) subtypes, and is also able to inhibit the growth of nPR(-) MB231 (TNBC-Caucasian American Women) cells [27–29]. Among TNBCs, over 70% are basal phenotype, one of the most aggressive TNBCs, and mPRs role in tumorigenesis, especially in basal phenotype, is of great interest since basal phenotype-derived breast cancer cells do not express nPRs [30].

In our previous studies, utilizing a multi-omics and systems biology approach, at both the transcriptional and translational levels, we identified 21 clinically relevant AAW-TNBC specific biomarkers with differential expression, in basal A (BaA) MDA-MB-468 (MB468) cells with significant survival curves, generating a collection of intrinsic and mPRs-specific steroid inducible biomarkers for AAW-TNBCs [31]. In this study, utilizing similar techniques, we are extending our previous findings by utilizing basal phenotype breast cancer line basal B (BaB) MDA-MB-231(MB231) cells, which also only express mPRs [32–34], to investigate the roles of key players of our previously defined CmP network [31] upon mPRs-specific steroid actions specifically in CAW-TNBCs. Our main objective is to understand the molecular regulatory mechanisms in breast cancer tumorigenesis under mPRs-specific steroid actions through cellular and molecular manipulation of CAW-TNBC cell lines *in-vitro*. Twenty-one potential biomarkers, associated with altered expression in CAW-TNBC cells/tissues, at both the transcriptional and translational levels, have been identified through our systems biology approach, which is significant for establishing a solid foundation for future CAW-TNBC therapeutic strategies.

## Results

### Dissecting the CmP network in CAW-derived TNBC cells using omics approaches

To dissect key players involved in the CmP signaling network, investigate key altered pathways affected in CAW-TNBC cells (MB231) under mPRs-specific steroid actions, and identify potential biomarkers, we examined the expression patterns at both the transcriptional and translational levels using high throughput RNAseq and LC-MS/MS omic approaches. In our previous findings utilizing AAW-TNBC cells (MB468) we observed that expression levels of key factors in the CmP signaling network differed between CAW and AAW TNBC cells, in which mPRs were enhanced at both the transcriptional and translational levels in Type 2A cells (CAW-TNBC) under mPRs- specific steroid actions, while mPRs expression was variable at both the transcriptional and translational levels in Type-2B cells (AAW-TNBC), suggesting multiple functions of the CmP signaling network for different TNBC subtypes [31]. As a result of these previous findings, we filtered out our RNAseq and proteomics data for CAW-TNBC cells by removing similarly altered Differentially Expressed Genes/Proteins (DEGs/DEPs) shared between Type 2B (AAW-TNBC) and Type 2A cells (CAW-TNBC) to evaluate CAW-TNBC specific DEGs/DEPs.

### Key signaling cascades identified within the CmP network in CAW-TNBC cells using RNAseq

Using the identified DEGs (Suppl. Table. 1), we performed hierarchical clustering to visualize changed gene expression between vehicle and mPRs-specific steroid treated CAW-TNBC cells (Fig. 1A). Further bioinformatics processing allowed us to visualize 200+ genes up-regulated and 350+ genes down-regulated, under mPRs-specific steroid actions, yielding a great foundation of potential biomarkers for type-2A TNBCs (Fig. 1B). These results also illustrated that in general, combined steroid actions have a stronger repressive role in the expression of CAW-TNBC specific DEGs (Fig. 1B). Pathway functional enrichment performed with these DEGs revealed several major altered signaling biological processes including overall down-regulation of cellular and metabolic, developmental, immune system as well as reproduction/reproductive processes in CAW-TNBC cells (Fig. 1C). Analysis of molecular functions affected revealed that the majority of alterations occurring in CAW-TNBC cells, under mPRs-specific steroid actions, are associated with an overall down-regulation in binding, catalytic activity as well as molecular function regulation pathways (Fig. 1D). Alteration of KEGG signaling pathways demonstrated that Wnt signaling was the major signaling pathway affected (Fig. 1E), which has major impacts on regulating development and stemness, and is involved in carcinogenesis in several major cancers including gastrointestinal [35, 36], leukemia [37–39], melanoma [40–43], and all sub-types of breast cancer [44, 45]. Additionally, Wnt signaling has been shown to affect the maintenance of cancer stem cells, metastasis, and immune control [46]. The other top signaling pathway altered in CAW-TNBC cells, under mPRs-specific steroid actions, included Inflammation mediated by chemokines and cytokines (Fig. 1E), which is known to contribute to tumor progression, in multiple cancers, specifically through mechanisms involved in migration, invasion, and metastasis [47]. Alterations in the Gonadotropin-releasing hormone (GnRH) signaling pathway (Fig. 1E) was also one of the top pathways affected, which is responsible for regulating mammalian reproduction, including production of hormones in both men and women [48]. Several angiogenesis-related pathways, heterotrimeric G-protein as well as integrin signaling pathways (Fig. 1E) were also among the top signaling pathways perturbed in CAW-TNBC cells. Overall, observations of altered key signaling cascades associated with tumorigenesis in CAW-TNBC cells, under mPRs-specific steroid actions, including apoptosis, p53, Wnt, RAS, CCKR, inflammation, angiogenesis, PDGF, and hormone signaling pathways (Fig. 1E), validate crosstalk between the CSC and mPRs in the CmP signaling network involved in CAW-TNBC tumorigenesis, and further validate the involvement of the CmP network in CAW-TNBC progression. Finally, to understand the classes of proteins primarily responsible for the observed alterations in key tumorigenesis signaling cascades, we performed further pathway functional enrichment which demonstrated the top 5 altered classes of proteins included metabolite interconversion enzymes, protein modifying enzymes, transporters, gene-specific transcriptional regulators, as well as nucleic acid metabolism proteins (Fig. 1F). Other classes of proteins contributing towards the observed altered pathways included protein binding activity modulators, transmembrane signal receptors, cytoskeletal proteins, cell adhesion, immunity, and structural proteins (Fig. 1F) in CAW-TNBC cells.

**Figure. 1.**
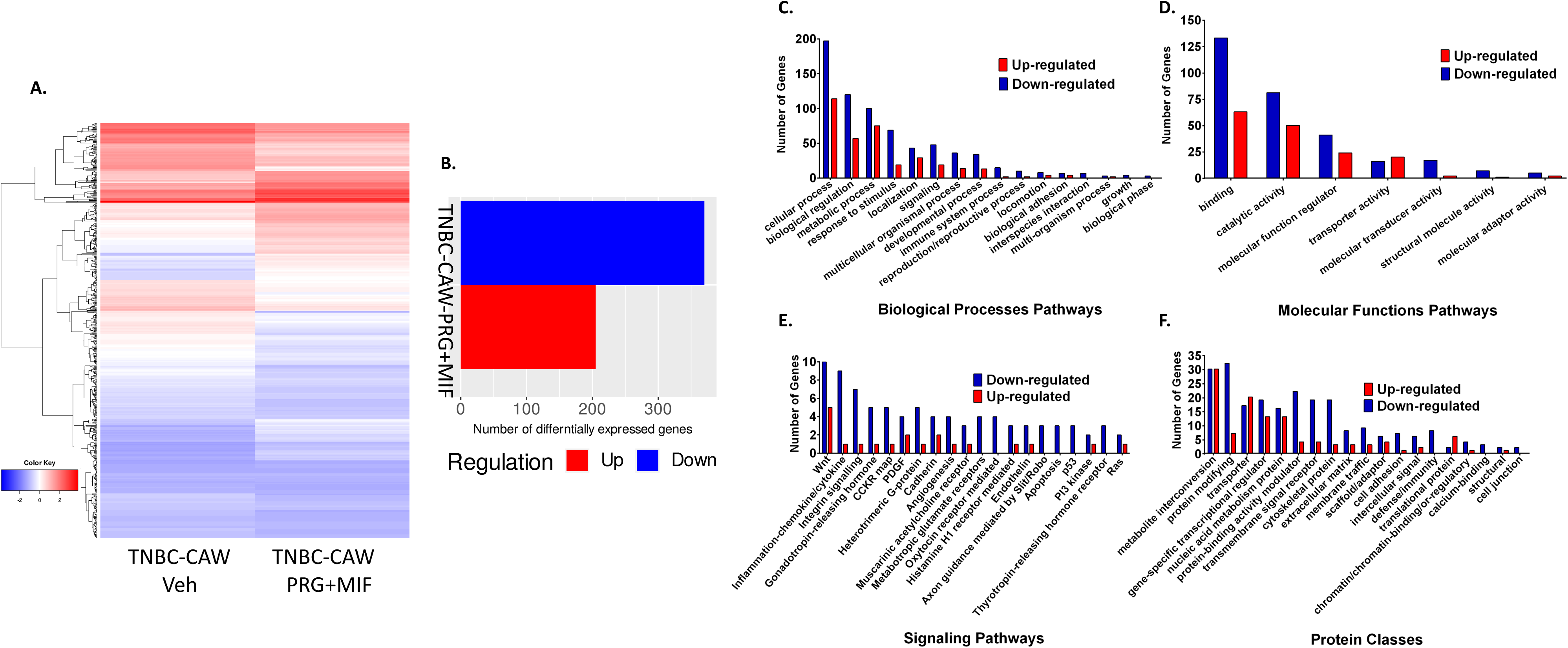
Differentially Expressed Genes (DEGs) among CAW-TNBC cells utilizing high-throughput RNA sequencing (RNAseq): CAW-TNBC (MB231) cells were treated with mPR specific steroids (MIF+PRG, 20µM each) for 48 hrs. Extracted significant DEGs, compared to vehicle controls, were further filtered using corresponding MB468 cells to obtain DEGs specific to CAW-TNBC cells. **A)** Cluster software and Euclidean distance matrixes were used for the hierarchical clustering analysis of the expressed genes (RNA) and sample program at the same time to generate the overall heatmap of changed gene expression. X-axis represents each comparing sample and Y-axis represents DEGs. **B)** Bioinformatics analysis was performed to identify significant DEGs, compared to vehicle controls. **C-F)** Pathway functional enrichment results for significant DEGs in mPR specific steroid MB231 treated cells, compared to their respective vehicle controls, were compiled using biological processes **(C)**, molecular functions **(D)**, KEGG signaling pathways **(E)**, and protein classes **(F)** data retrieved from the PANTHER (Protein ANalysis THrough Evolutionary Relationships) classification system. Additionally, all functional enrichment results were further stratified to identify up/down-regulation within each signaling pathway using iDEP (Integrated Differential Expression and Pathway Analysis) program. Coloring indicates the log2 transformed fold change (high: red, low: blue).

### Key signaling cascades within the CmP network in CAW-TNBC cells using LC-MS/MS approaches

Among the identified DEPs (Suppl. Table. 2), we again performed hierarchical clustering to visualize changed protein expression between vehicle and mPRs-specific steroid treated CAW-TNBC cells (Fig. 2A). Further bioinformatics processing demonstrated 50+ proteins up-regulated and 100+ proteins down-regulated, under mPRs-specific steroid actions (Fig. 2B). These results, in complementation to potential DEG biomarkers identified (Fig. 1B), provided us with an additional index of potential CAW-TNBC specific DEP biomarkers to be further investigated. These results also illustrated that in general, mPRs-specific steroid actions have a stronger repressive role in the expression of CAW-TNBC specific DEPs (Fig. 2B). Protein functional enrichment demonstrated that several pathways overlaid with RNA enrichment, reinforcing that these pathways may play potential key roles in hormone signaling and factors in CAW-TNBCs (Figs. 1C-F and 2C-F). Pathway functional enrichment performed with DEPs revealed all the same major altered signaling biological processes seen at the transcriptional level, including overall down-regulation of cellular, metabolic, developmental, immune system as well as reproduction/reproductive processes in CAW-TNBC cells (Fig. 2C). Similarly, analysis of molecular functions also revealed that the majority of alterations, identical to the results obtained at the transcriptional level, are associated with an overall down-regulation in binding, catalytic activity, structural molecule activity, as well as molecular function regulation pathways (Fig. 2D). Surprisingly, Wnt was not the most impacted KEGG pathway at the translational level, as was seen at the transcriptional level, but instead down-regulation of angiogenesis was the most impacted signaling pathway in mPRs-specific steroid treated CAW-TNBC cells (Fig. 2E), which was also in the top affected signaling pathways observed in our RNAseq data (Fig. 1E). At the translational level, Wnt signaling was still in the top 15 pathways disrupted, reinforcing its definitive role in CAW-TNBC cells. Synonymous with our transcriptomics data, the top signaling pathway altered in CAW-TNBC cells, under mPRs- specific steroid actions, included Inflammation mediated by chemokines and cytokines, GnRH, heterotrimeric G-protein, integrin, apoptosis, p53, RAS, CCKR, and PDGF signaling pathways (Fig. 2E). These results further reinforce the existence of crosstalk between the CSC and mPRs in the CmP signaling network involved in CAW-TNBC tumorigenesis, and further validate the involvement of the CmP network in CAW-TNBC progression. It must be noted that overall, the number of proteins identified for each pathway in our proteomics study is greatly decreased from the number of genes identified in our RNAseq analysis, which is not surprising given the reduced sensitivity of proteomics compared to RNAseq, however, the identification of the same top affected signaling pathways validates the results seen at both levels independently. Similar to our transcriptomics analysis, 3 of the top 5 altered classes of proteins included metabolite interconversion enzymes, protein modifying enzymes, and nucleic acid metabolism proteins with the other top two including cytoskeletal, and translational proteins (Fig. 2F) in mPRs-specific steroid treated CAW-TNBC cells. Together, these results solidify the existence of a cellular relationship between the CSC and mPRs signaling, at both the transcriptional and translational levels, in CAW-TNBC cells. In addition to identifying key perturbed signaling pathways, our extensive omics analysis has also provided several new candidate biomarkers that can be further investigated to determine their potential clinical applications for CAW-TNBCs.

**Fig. 2.**
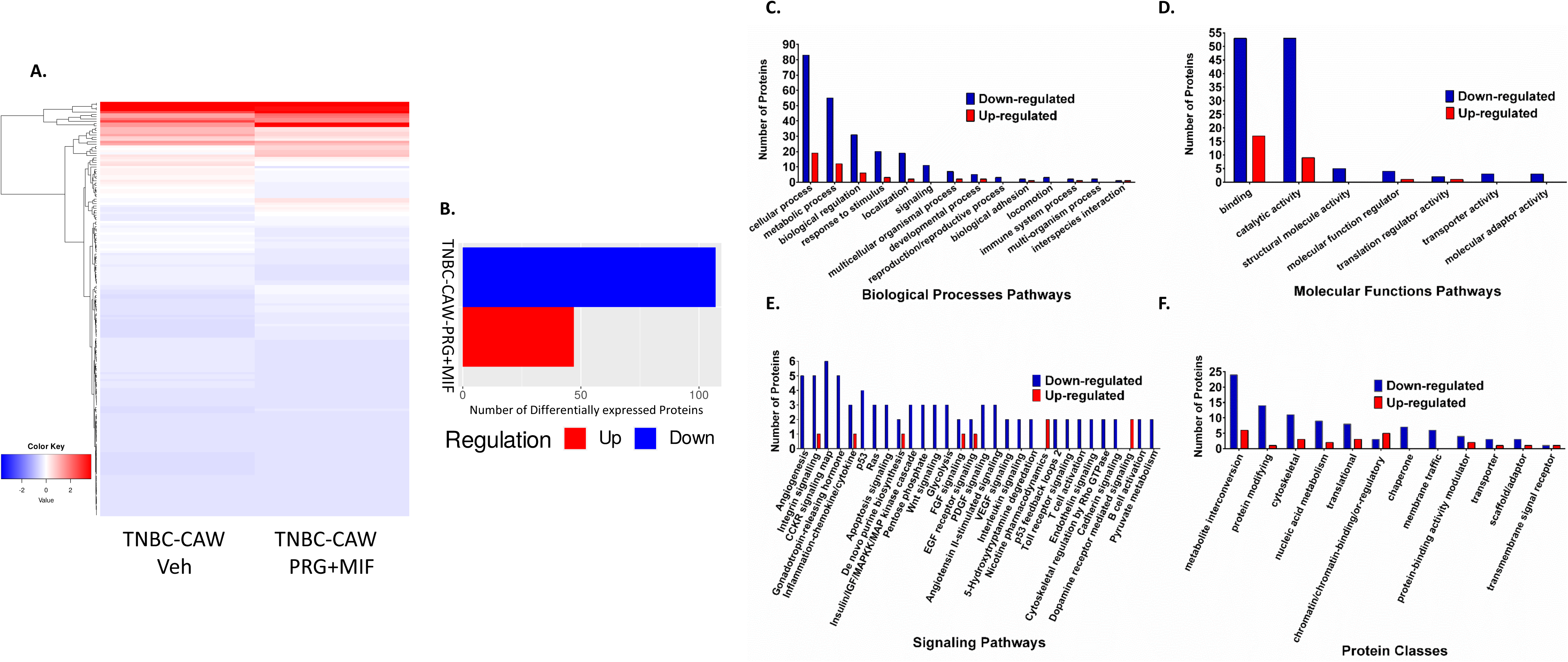
Differentially Expressed Proteins (DEPs) among CAW-TNBC cells utilizing high-throughput Proteomics: CAW-TNBC cells (MB231) were treated with mPR specific steroids (MIF+PRG, 20µM each) for 48 hrs. Extracted significant DEPs, compared to vehicle controls, were further filtered using corresponding MB468 cells to obtain DEPs specific to CAW-TNBC cells. **A)** Cluster software and Euclidean distance matrixes were used for the hierarchical clustering analysis of the expressed proteins and sample program at the same time to generate the overall heatmap of changed protein expression using *t-test* statistical analysis. **B)** Bioinformatics analysis was performed to identify significant DEPs, compared to vehicle controls. **C-F)** Pathway functional enrichment results for significant DEPs in mPR specific steroids treated MB231 cells, compared to their respective vehicle controls, were compiled using biological processes **(C)**, molecular functions **(D)**, KEGG signaling pathways **(E)**, and protein classes **(F)** data retrieved from the PANTHER (Protein ANalysis THrough Evolutionary Relationships) classification system. Additionally, all functional enrichment results were further stratified to identify up/down-regulation within each signaling pathway using iDEP (Integrated Differential Expression and Pathway Analysis) program. Coloring indicates the log2 transformed fold change (high: red, low: blue).

### Key signaling cascades within the CmP network in CAW-TNBC cells identified using systems biology approaches

To evaluate the inter-relationship between RNA and protein expression from CAW-TNBC cells, under mPRs-specific steroid actions, we overlaid the regulation patterns and extrapolated the overlaps between the two datasets producing a list of 32 CAW-TNBC specific potential biomarkers that were seen perturbed at both the transcriptional and translational levels (Fig. 3A, Suppl. Table 3). Overall down-regulation of biological processes, molecular functions and protein classes affected with these overlapped DEGs/DEPs (Figs. 3B, 3C, 3E) were nearly identical to the individual analysis seen. Down-regulation of 18 KEGG pathways, were also very similar to the individual analysis performed with the RNAseq and proteomics data, however, the top affected signaling pathway using our reduced list of potential biomarkers demonstrated P53 signaling at the top, followed by integrin, inflammation by chemokines and cytokines, Wnt and apoptosis signaling, all seen at the individual levels as well (Fig. 3D). Other consistently down-regulated pathways also identified in the individual analysis included angiogenesis, GnRH signaling, as well as heterotrimeric G-protein pathways (Fig. 3D), further validating our independent observations at both the transcriptional and translational levels. To reduce the list of identified potential biomarkers, we reviewed expression patterns between DEGs/DEPs (Fig. 3A), and only genes/proteins with the same regulation trends (averaged log2 Fold Changes) seen at both the transcriptional and translational levels were further assessed (29 DEGs/DEPs). Genes/proteins displaying opposite expression at the transcriptional and translational levels (3 DEGs/DEPs) were excluded from further analysis as these reflected genes capable of undergoing potential feedback auto-regulation in CAW-TNBC cells under mPRs-specific steroid actions.

**Figure 3.**
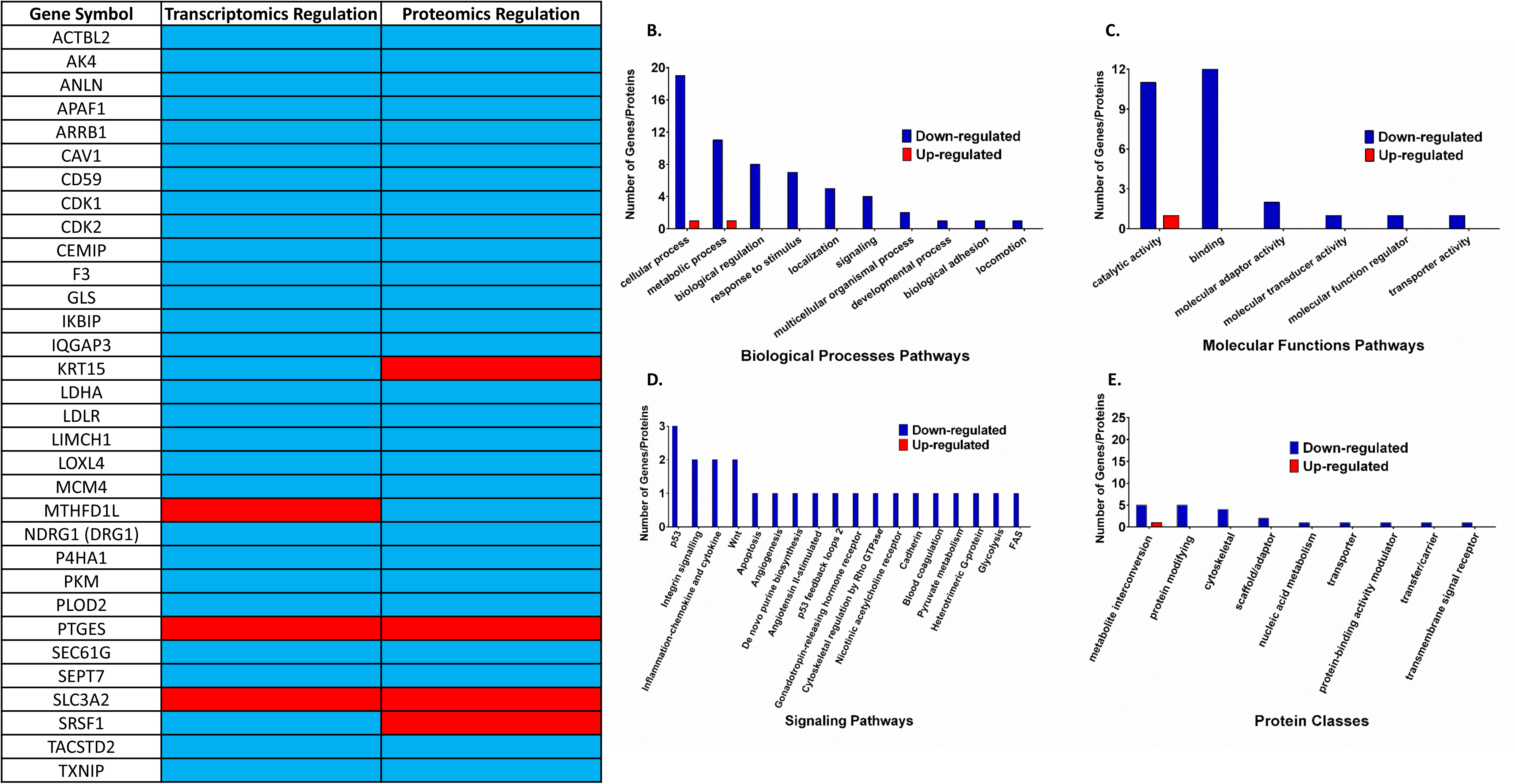
Overlapped Differentially Expressed Genes/Proteins (DEGs/DEPs) among CAW-TNBC cells utilizing high-throughput RNA sequencing (RNAseq) and Proteomics. CAW-TNBC cells (MB231) were treated with mPR specific steroids (MIF+PRG, 20µM each) for 48 hrs. Overlapped extracted significant DEGs/DEPs, compared to vehicle controls, were further filtered using corresponding overlapped extracted significant DEGs/DEPs in MB468 cells to obtain 32 DEGs/DEPs specific to CAW-TNBC cells to serve as the potential list of candidate CAW-TNBC biomarkers. **(A)** 32 overlapped significant DEGs/DEPs were identified in our comparative omics analysis, and are displayed with their corresponding regulation at both the transcriptional and proteomic levels. **B-E)** Pathway functional enrichment results for overlapped significant DEGs/DEPs in mPR specific steroids treated MB231 cells, compared to their respective vehicle controls, were compiled using biological processes **(B)**, molecular functions **(C)**, KEGG signaling pathways **(D)**, and protein classes **(E)** data retrieved from the PANTHER (Protein ANalysis THrough Evolutionary Relationships) classification system. Additionally, all functional enrichment results were further stratified to identify up/down-regulation within each signaling pathway using iDEP (Integrated Differential Expression and Pathway Analysis) program. Coloring indicates the log2 transformed fold change (high: red, low: blue).

### Differential expression of our candidate biomarkers and their associated prognostic effects

We next set out to filter out our 29 remaining candidate biomarkers for CAW-TNBCs from our multi-omics analysis to evaluate their clinical relevance. To accomplish this, we analyzed publicly available microarray data [49] for the remaining 29 candidate biomarkers to evaluate prognostic effects by integrating gene expression and clinical data simultaneously to generate Kaplan-Meier (KM) survival curves. Breast cancer patient samples were filtered to only analyze patient samples classified as ER(-)/nPR(-)/HER2(-)/TNBC subtype. For the first group of biomarkers, a worst prognostic effect was illustrated corresponding with our observed differential expression in CAW-TNBCs (Figs. 3A), under mPRs-specific steroid actions, for SLC3A2, IKBIP, ARRB1, CD59, GLS, LDLR, MCM4, PTGES, and TXNIP (Suppl. Fig. 1A-C). These results suggest that a disrupted CmP signaling network, under mPRs-specific steroid actions, in TNBC cells could lead to worst prognostic outcomes. Conversely, for the second group of biomarkers, a better prognostic effect with our observed differential expression in CAW-TNBCs (Figs. 3A), under mPRs-specific steroid actions, for AK4, CAV1, F3, TACSTD2, LDHA, PLOD2, APAF1, CDK2, IQGAP3, DRG1 (NDRG1), P4HA1, and SEPT7 was observed (Suppl. Fig. 1D-F). Together these results demonstrate the heterogeneity response of CAW-TNBCs under mPRs-specific steroid actions. For 5 biomarkers, we did not obtain significant clinical relevance with differential expression in TNBCs (Suppl. Fig. 2), and 3 of our initially identified biomarkers did not have any data available to perform the analysis. Utilizing our KM survival curve data (regardless of prognostic effect), we further assessed our biomarkers by only proceeding with biomarkers that had significant KM survival curves (21 biomarkers) in TNBC patients to assess their basal expression patterns in clinical samples.

### Investigating differential expression of our candidate biomarkers in breast cancer tissues

With our remaining 21 clinically significant CAW-TNBC specific candidate biomarkers, we next sought out to evaluate their differential expression among breast cancer subtypes. Utilizing publicly available breast cancer tumor tissue gene expression data (microarray) [49] we explored differential expression patterns for our candidate biomarkers, associated with significant KM curves identified in this study, in nPR(+/-) breast cancer tissues (ER and HER2 receptor status not available). Expression data, from breast cancer tissue microarrays, were divided based on Progesterone Receptor status (nPR+/-). 12 of our candidate biomarkers, AK4, APAF1, CD59, CDK2, LDHA, GLS, IQGAP3, PLOD2, NDRG1, SLC3A2, P4HA1, and TACSTD2 were found to be significantly up-regulated in nPR(-) tumor tissues, compared to nPR(+) tumor tissues (Suppl. Fig.3A-C). Interestingly, we only observed opposite up-regulation for 1/12 biomarkers (SLC3A2) in CAW-TNBC cells under mPRs-specific steroid actions (Fig. 3A), suggesting little effect of hormone actions on SLC3A2 expression levels. The other 11 biomarkers, however, were down-regulated in CAW-TNBC cells, under mPRs-specific steroid actions (Fig. 3A), suggesting that disruption of the CmP signaling network has a huge influence in altering expression for these genes/proteins at both the transcriptional and translational levels. Alternatively, 4 of our candidate biomarkers, ARRB1, F3, PTGES, and TXNIP were found to be significantly down-regulated in nPR(-) compared to nPR(+) tumor tissues (Suppl. Fig. 3D). Interestingly, we observed opposing up-regulation for 1/4 biomarkers (PTGES) in CAW-TNBC cells (Fig. 3A), under mPRs-specific steroid actions, suggesting that disruption of the CmP signaling network has a huge influence in altering expression for this naturally under-expressed gene in nPR(-) tumor tissues. The other 3 biomarkers (ARRB1, F3, and TXNIP) demonstrated similar down-regulated patterns in our omics data (Fig. 3A), suggesting little effect of hormone actions on their expression levels. Furthermore, 5 biomarkers, CAV1, IKBIP, LDLR, MCM4, and SEPT7 were found to have similar expression patterns in nPR(+/-) tumor tissues (Suppl. Fig. 4). However, disruption of the CmP signaling network, under mPRs-specific steroid actions in CAW-TNBC cells resulted in significant down-regulation of all 5 biomarkers (Fig. 3A) suggesting that combined steroid actions induces decreased expression, at both the transcriptional and translational levels for these 5 biomarkers in CAW-TNBC cells.

We next sought to further our analysis to investigate differential expression patterns between AAW-TNBCs and CAW-TNBCs utilizing publicly available RNA-seq data comparing expression levels in 23 AAW-TNBC and 19 CAW-TNBC samples [50] for our candidate biomarkers associated with significant KM curves identified in this study. ARRB1, CDK2, IQGAP3, MCM4, NDRG1, and TACSTD2 all displayed similar down-regulated trends in CAW-TNBC cells (Fig. 3A), under mPRs-specific steroid actions, as was naturally observed in the CAW-TNBC patient samples (Suppl. Fig 3E), allowing us to define these genes/proteins as down-regulated intrinsic biomarkers for CAW-TNBCs (Fig. 4A, blue-colored biomarkers, column 1). Two representative examples of KM survival curves (Fig. 4B) and differential expression patterns in CAW tissues (Fig. 4C) corresponding to our observed omics analysis (Fig. 3A) is illustrated for 2 intrinsic biomarkers. Additionally, PTGES which displayed similar up-regulated trends in CAW-TNBC cells (Fig. 3A), under mPRs-specific steroid actions, as was naturally observed in the CAW-TNBC patient samples (Fig. 4C and Suppl. Fig. 3F-panel viii) was the only up-regulated intrinsic biomarker identified in our study (Fig. 4A, red-colored biomarker, column 1). Alternatively, AK4, CAV1, APAF1, F3, CD59, LDHA, GLS, PLOD2, LDLR, SLC3A2, P4HA1, SEPT7, IKBIP, and TXNIP displayed either opposite or inducible trends in CAW-TNBC cells (Fig. 3A), under mPRs-specific steroid actions, as was naturally observed in the CAW-TNBC patient samples, (Suppl. Fig. 3F all panels except panel viii, and Suppl. Fig. 5), allowing us to define these genes/proteins as inducible biomarkers (CmP disruption) for CAW-TNBCs (Fig. 5A, black colored biomarkers, column 1). Representative examples of KM survival curves (Fig. 5B) and differential expression patterns in CAW tissues (Fig. 5C) corresponding to our observed omics analysis (Fig. 3A) is illustrated for 2/14 inducible biomarkers. Utilizing all of the expression and prognostic data, we were able to filter our initial 32 candidate biomarker hits (Fig. 3A and Suppl. Table 3) to a condensed, clinically relevant list of 21 intrinsic and inducible biomarkers for CAW-TNBCs (Table 1).

**Fig. 4.**
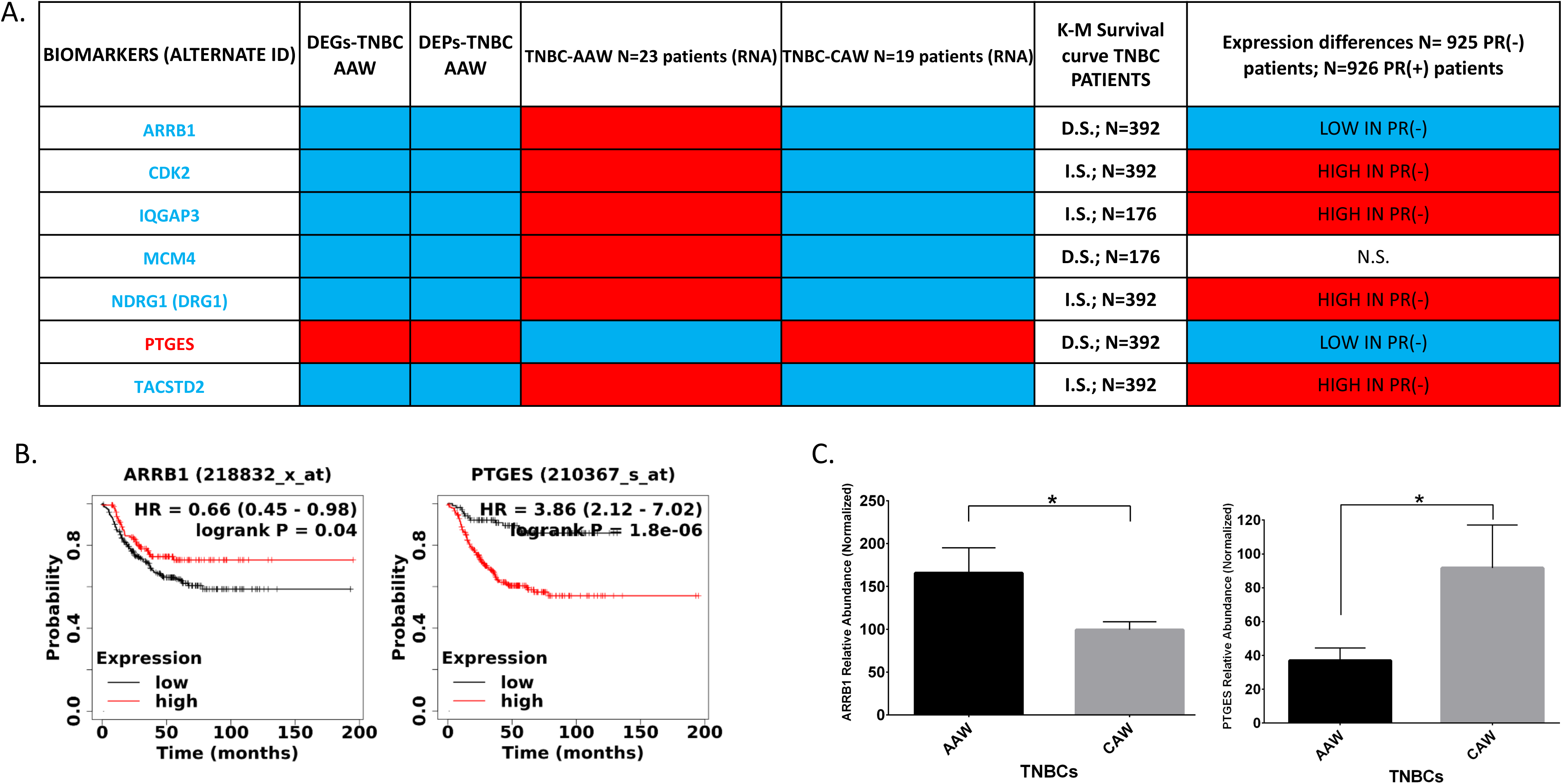
Intrinsic biomarkers identified among CAW-TNBC cells utilizing high-throughput RNA sequencing (RNAseq) and Proteomics. Overlapped synchronous DEGs/DEPs in CAW-TNBCs identified as intrinsic biomarkers. **A)** Systems biology data for the 7 intrinsic biomarkers identified in this study are summarized which evaluated **B)** a set of publicly available microarray data (22,277 probes) from 1,809 breast cancer samples using kmplot software to integrate gene expression and clinical data simultaneously to generate the displayed Kaplan-Meier survival curve results which demonstrated significant prognosis effects with our observed results (2 representative figures illustrated), as well as **C)** assessing expression levels in AAW-TNBC (n=23) and CAW-TNBC (n=19) tumor tissues which all displayed similar trends with a disrupted CmP signaling network, as was naturally observed in the CAW-TNBC patient samples (2 representative figures illustrated), allowing us to define these genes/proteins as intrinsic biomarkers for CAW-TNBCs.

**Fig. 5.**
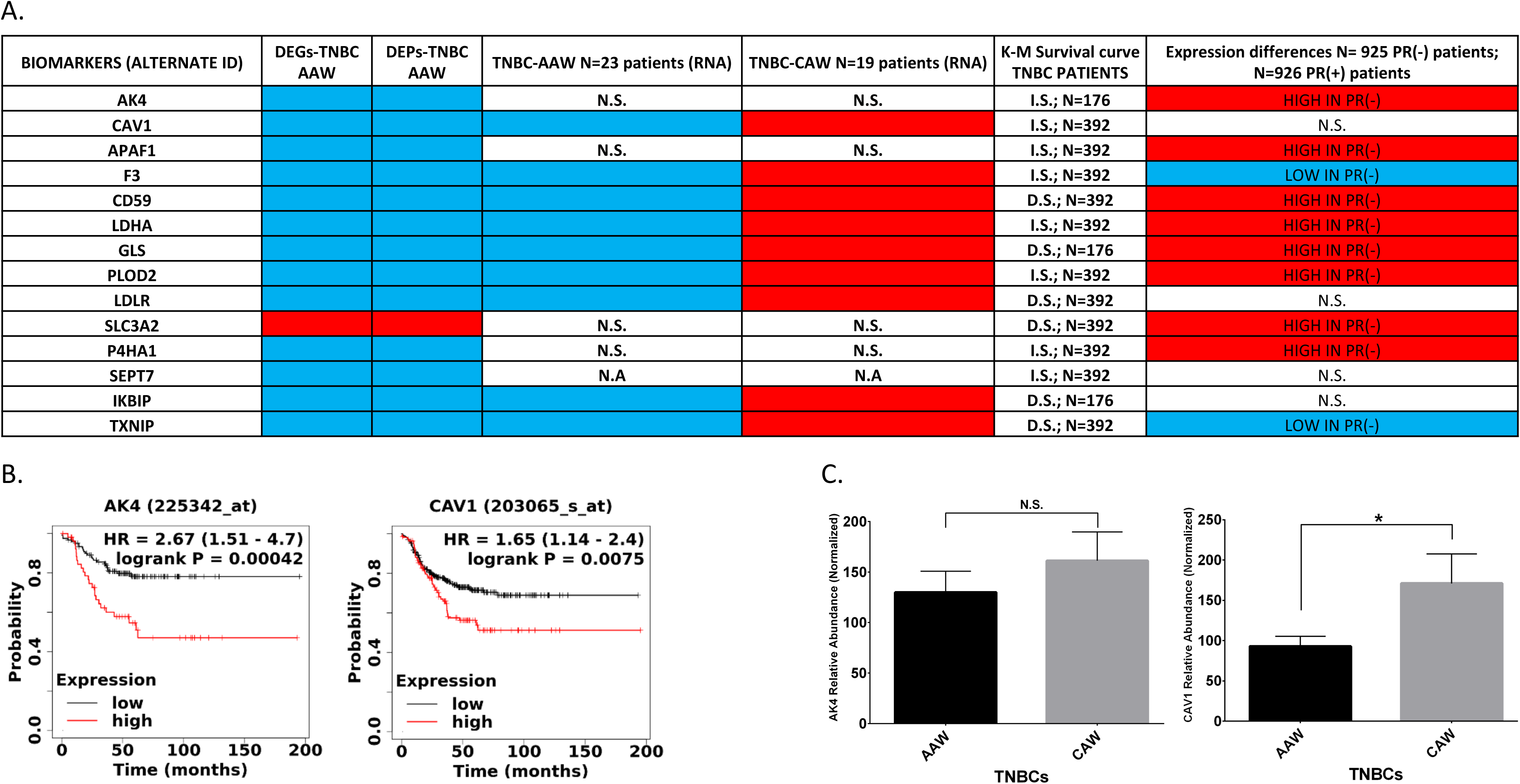
Inducible biomarkers identified among CAW-TNBC cells utilizing high-throughput RNA sequencing (RNAseq) and Proteomics. Overlapped synchronous DEGs/DEPs in CAW-TNBCs identified as inducible biomarkers. **A)** Systems biology data for the 14 intrinsic biomarkers identified in this study are summarized which evaluated **B)** a set of publicly available microarray data (22,277 probes) from 1,809 breast cancer samples using kmplot software to integrate gene expression and clinical data simultaneously to generate the displayed Kaplan-Meier survival curve results which demonstrated significant prognosis effects with our observed results (2 representative figures illustrated), as well as **C)** assessing expression levels in AAW-TNBC (n=23) and CAW-TNBC (n=19) tumor tissues which all displayed opposite or inducible trends with a disrupted CmP signaling network, as was naturally observed in the CAW-TNBC patient samples (2 representative figures illustrated), allowing us to define these genes/proteins as inducible biomarkers for CAW-TNBCs.

**Table 1.** Summarized 21 identified prognostic candidate biomarkers associated with clinically significant data in CAW-TNBCs. Candidate biomarkers shared at both RNA/Protein levels under hormone treatments (columns 1, 2, 3), were further validated in AAW-TNBC (n=23) and CAW-TNBC (n=19) (columns 4, 5) tumor tissues. These biomarkers were further analyzed by a set of publicly available microarray data (22,277 probes) from 1,809 breast cancer samples using kmplot software to integrate gene expression and clinical data simultaneously to generate the displayed Kaplan-Meier survival curve results (column 6). Candidate biomarkers were further assessed with two groups of breast cancer tissue samples, based on classic progesterone Receptor (nPR) status determined by Immunohistochemistry (IHC), to evaluate clinical expression levels focusing on 436-925 nPR(-) (depending on the probe) and 511-926 nPR(+) (depending on probe) breast cancer samples (column 7). Red-colored background indicates increased/higher expression levels, while blue-colored background indicates decreased/lower expression levels. Colored biomarkers (red or blue in Column 1) indicate intrinsic biomarkers which are selected as candidate biomarkers for AAW-TNBCs, while black colored biomarkers are inducible biomarkers. Abbreviations: I.S., Increased Survival; D.S., Decreased Survival; N.S., No statistical significance; HIGH IN nPR(-), Higher expression levels in PR(-) breast cancer tissues; LOW IN PR(-), Lower expression levels in nPR(-) breast cancer tissues; N/A, Not Available.

| BIOMARKERS (ALTERNATE ID) | DEGs-TNBC<br>AAW | DEPs-TNBC<br>AAW | TNBC-AAW N=23 patients (RNA) | TNBC-CAW N=19 patients (RNA) | K-M Survival<br>curve TNBC<br>PATIENTS | Expression differences N= 925 PR(-) patients;<br>N=926 PR(+) patients |
| --- | --- | --- | --- | --- | --- | --- |
| AK4 |  |  | N.S. | N.S. | I.S.; N=176 | HIGH IN PR(-) |
| CAV1 |  |  |  |  | I.S.; N=392 | N.S. |
| APAF1 |  |  | N.S. | N.S. | I.S.; N=392 | HIGH IN PR(-) |
| ARRB1 |  |  |  |  | D.S.; N=392 | LOW IN PR(-) |
| F3 |  |  |  |  | I.S.; N=392 | LOW IN PR(-) |
| CD59 |  |  |  |  | D.S.; N=392 | HIGH IN PR(-) |
| CDK2 |  |  |  |  | I.S.; N=392 | HIGH IN PR(-) |
| LDHA |  |  |  |  | I.S.; N=392 | HIGH IN PR(-) |
| GLS |  |  |  |  | D.S.; N=176 | HIGH IN PR(-) |
| IQGAP3 |  |  |  |  | I.S.; N=176 | HIGH IN PR(-) |
| PLOD2 |  |  |  |  | I.S.; N=392 | HIGH IN PR(-) |
| LDLR |  |  |  |  | D.S.; N=392 | N.S. |
| MCM4 |  |  |  |  | D.S.; N=176 | N.S. |
| NDRG1 (DRG1) |  |  |  |  | I.S.; N=392 | HIGH IN PR(-) |
| SLC3A2 |  |  | N.S. | N.S. | D.S.; N=392 | HIGH IN PR(-) |
| P4HA1 |  |  | N.S. | N.S. | I.S.; N=392 | HIGH IN PR(-) |
| PTGES |  |  |  |  | D.S.; N=392 | LOW IN PR(-) |
| SEPT7 |  |  | N.A | N.A | I.S.; N=392 | N.S. |
| IKBIP |  |  |  |  | D.S.; N=176 | N.S. |
| TXNIP |  |  |  |  | D.S.; N=392 | LOW IN PR(-) |
| TACSTD2 |  |  |  |  | I.S.; N=392 | HIGH IN PR(-) |

### Differential expression of shared TNBC candidate biomarkers and their associated prognostic effects

In an effort to identify DEGs/DEPs shared between AAW-TNBCs and CAW-TNBCs, we overlapped omics data between the two datasets (unfiltered) to identify shared synchronous DEGs/DEPs to enhance our biomarker analysis. Interestingly, our analysis only resulted in 2 shared potential biomarkers between the two TNBC cell lines (Suppl. Fig 6). Our systems biology analysis revealed that PCNA is a potential intrinsic biomarker for TNBCs with a significant KM survival curve (Suppl. Fig. 6A), however, we did not obtain significant differential expression in either nPR(+/-) tissues (Suppl. Fig. 6B) or between AAW-TNBCs and CAW-TNBCs (Suppl. Fig. 6C). Our second identified biomarker, SRSF2, appears to undergo potential feedback auto-regulation in both AAW-TNBC and CAW-TNBC cells under mPRs-specific steroid actions (Suppl. Fig. 6D), eliminating it as a potential biomarker in TNBCs.

## Discussion

The existence of the PRG-mPRs signaling cascade in nPR(+/-) cells have been well documented [30, 33, 51–53], demonstrating that PRG signaling in nPR(-) cell lines is solely mediated through mPRs (PAQRs) [30]. In our previous study, we defined the novel CmP signaling network in TNBC cells [31], which overlapped with our previously defined CmPn network in nPR(+) breast cancer cells [25], and observed that combined steroid actions have a positive effect on CSC protein expression in TNBCs with inducible patterns of expression in AAW-TNBC cells [31]. Using extensive multi-omics approaches, we were once again able to demonstrate alterations to key tumorigenesis pathways including cytokine-mediated responses, Wnt, Integrin, GnRH, and angiogenesis signaling pathways in CAW-TNBC cells under mPRs-specific steroid actions, which were in parallel to the observation in AAW-TNBC cells under identical conditions [31]. These results suggest that even though TNBC diagnoses in AAW are associated with more aggressive forms of the disease [54–56], and experience a higher mortality rate [57], TNBC in CAW share similar altered signaling pathways, under mPRs-specific steroid actions, demonstrating the aggressive nature of TNBCs regardless of racial differences. Utilizing a similar multi-omics approach, we identified 21 new CAW-TNBC specific candidate biomarkers (Table 1) that will be further validated to evaluate their potential clinical applications for CAW-TNBCs utilizing patient derived xenograft (PDX) mouse models. In addition, this project reinforces the definitive role of the CmP signaling network in TNBC tumorigenesis that was initially identified in our previous studies with AAW-TNBCs [31]. This new set of potential prognostic biomarkers for CAW diagnosed with TNBC may revolutionize molecular mechanisms and currently known concepts of tumorigenesis in CAW-TNBCs, leading to hopeful new therapeutic strategies.

### Potential prognostic biomarkers for CAW-TNBCs

Biomarkers, identified through cutting-edge biomedical technologies (such as genetics, epigenetics, genomics, proteomics, etc.), are prognostic tools that can be used to predict patient response to therapy and future course of the disease [58], and have been widely used in breast cancer therapies to offer precise treatment based on tumor characteristics [59]. As mentioned previously, TNBC patients have limited options for treatment since most immunotherapies target the ER, nPR, and HER2 receptors, which are ineffective for TNBCs [19]. TNBCs, among all other breast cancer subtypes, have the most limited amount of therapeutic strategies available, demonstrate limited responses to currently available treatments, and have the worst prognosis [10, 13, 19, 20]. Therefore, the identification of new biomarkers for TNBC subtypes is of the utmost importance in guiding treatment decisions and predicting prognosis in the future for precision-based cancer therapies [60–66].

Utilizing our expertise in multi-omics analysis, we utilized CAW-derived TNBC cells to discover a total of 29 potential biomarkers synchronously regulated at the transcriptional and translational levels, associated with tumorigenesis in CAW-derived TNBCs (Fig. 3A). Furthermore, utilizing our expertise in systems biology analysis, we utilized publicly available clinical data (expression and prognosis), to filter the 29 biomarkers down to 21 clinically relevant CAW-TNBC specific biomarkers demonstrating differential expression, under mPRs-specific steroid actions, and demonstrating significant survival curves (Table 1). This analysis resulted in the identification of 7 intrinsic and 14 inducible biomarkers for CAW-TNBCs (Table 1). Despite previous reports of DEGs/DEPs between TNBC basal subtypes [67–70], our results demonstrated that disruption of the CmP signaling network, through mPRs-specific steroid actions in CAW-TNBCs, induces differential expression patterns of key players in the CmP network, as well as disrupts key tumorigenesis signaling pathways (Figs. 1-3). Furthermore, our findings do not overlap with any previous publications and are unrelated to the reported DEGs/DEPs in TNBC subtypes, yielding our newly identified biomarkers as great candidates for CAW-TNBC clinical applications (Table 1).

### Pathways functional enrichment analysis using DEGs/DEPs demonstrate perturbation in key tumorigenic signaling cascades

Pathways functional enrichment results obtained from significantly altered genes/proteins in CAW-TNBCs, under mPRs-specific steroid actions, demonstrated perturbation in key signaling cascades known to participate in tumorigenesis and cancer progression. One of the main altered pathways, seen at both the transcriptional and translational levels, included inflammation by chemokines and cytokines. These proteins are known to have a vast range of functionalities including preventing apoptosis thereby inducing proliferation of cancer cells [71] and contributing to induction/metastasis as well as homeostatic regulation of breast cancer [72]. Additionally, they are also suspected to modulate tumor growth by stimulating angiogenic factors in tumor cells [73]. Another key signaling pathway affected in CAW-TNBCs, under mPRs-specific steroid actions included Wnt which has been shown to affect the maintenance of cancer stem cells, metastasis, immune control, and is responsible for signal transduction through cell surface receptors [46]. Wnt, classified as a proto-oncogene, can induce uncontrolled cell division [74] and has been associated with various cancers including lung [75], prostate [76], gastrointestinal [35, 36], leukemia [37–39], melanoma [40–43], and all sub-types of breast cancer [44, 45].

Altered integrin signaling, was also observed in our multi-omics analysis in CAW-TNBCs, under mPRs-specific steroid actions, and has been shown to aid in cell proliferation, adhesion, growth, division, apoptosis, and is suspected to induce angiogenesis, another major altered pathway in our functional enrichment analysis, through activation of MAPKs [77]. Furthermore, regulation of Integrins, along with IGF-IR in mitogenesis, has been shown to contribute towards prostate cancer development [78]. Modulation of the GnRH pathway, another major pathway affected in our enrichment analysis with CAW-TNBCs under mPRs-specific steroid actions, is responsible for regulating mammalian reproduction, including production of hormones in both men and women [48] and is known to contribute towards endometrium, prostate, and breast cancer progression [79]. Interestingly, Integrins and GnRH, are known to mediate one another in ovarian cancer progression [80]. These observations further validate the possible existence of crosstalk between the CSC and mPRs, in the CmP signaling network involved in tumorigenesis pathways, and further validate the involvement of the CmP network in CAW-TNBC progression observed in this study.

### Intrinsic and inducible CAW-TNBC biomarkers have known roles in cancer progression

Identification of novel CAW-TNBC biomarkers with clinical relevance is of great importance, especially due to limited treatment options [19]. During our studies, we found an overwhelming amount of literature support, demonstrating that the majority of our identified biomarkers had confirmed roles in signaling pathways confirmed in cancer progression, validating their potential as prognosis biomarkers in CAW-TNBCs. AK4, observed to be down-regulated in our studies, is involved in the progression of Serous ovarian cancer [81], and AK4 depletion was shown to impair cell proliferation and invasion in MCF7 as well as MB231 breast cancer cells [82]. CAV1, a tumor suppressor gene candidate, is a negative regulator of the Ras-p42/44 mitogen-activated kinase cascade [83], and has been shown to play a critical role in the progression of breast cancer through autophagy, invasion, proliferation, apoptosis, migration, and breast cancer metastasis [84]. Furthermore, CAV1 has demonstrated involvement in radiotherapy and chemotherapy resistance, which are key issues encountered in breast cancer treatment, and stromal CAV1 has been proposed as a potential indicator for breast cancer prognosis [84]. APAF1, suppressed in our studies, is mutated in human melanomas, and its depletion has been shown to contribute towards malignant transformations in cancerous mouse models [85]. Over-expression of ARRB1, identified here as an intrinsic down-regulated biomarker, reduced TNBC cell migration and growth, and expression levels have been reported to inversely correlate with histological grade and positively associate with patient survival, suggesting a tumor-suppressive role for ARRB1 in TNBCs [86]. F3 has been shown to increase tumor growth through enhancing integrin β1 signaling and contributes to metastasis through thrombin mechanisms that protect tumor cells from natural killer cells [87]. CD59 over-expression promotes proliferation of MCF-7 breast cancer cells while simultaneously decreasing Bcl-2 expression, which is a strategy used by tumor cells to evade complement-dependent cell cytotoxicity stimulated by monoclonal antibodies [88, 89]. Interestingly, CDK2 has been shown to activate poly-(ADP)-ribose polymerase 1, which is required for hormonal gene regulation in breast cancer cells, and the connecting signaling pathway involved with this process has been proposed as a novel pathway for pharmacological management of breast cancer [90]. LDHA, down-regulated in our studies, is normally over-expressed in TNBCs, and expression of LDHA was significantly correlated with TNM staging, distant metastasis, and survival outcomes; its expression levels along with AMPK have been proposed as prognostic biomarkers in breast cancer [91]. GLS, involved in both breast and colorectal cancer, through its ability to deregulate glutaminolysis, and has been linked to cancer progression for both types [92, 93]. IQGAP3, proposed as a novel diagnostic/prognostic marker and therapeutic target for both pancreatic and serous ovarian cancer, is suspected to act as an oncogene by regulating Cdc42 expression in pancreatic cancer [94] and was shown to promote cancer proliferation and metastasis in high-grade serous ovarian cancer [95]. PLOD2, contributing towards drug resistance in laryngeal cancer [96] as well as regulating peritoneal dissemination in gastric cancer [97], has also been shown to be critical for fibrillary collagen formation in breast cancer cells, contributing to tumor stiffness, and the main mechanism for metastasis to the lungs and lymph nodes [98]. MCM4 expression has been associated with the pathological staging of esophageal cancer [99] and has also been shown to play an essential role in the proliferation of non-small cell lung cancer [100]. NDRG1 and its interaction with Cap43 have been shown to predict tumor angiogenesis and has been proposed as a poor prognosis marker in non-small-cell lung cancer using *in-vivo* animal models [101]. Furthermore, NDRG1 expression has been associated with breast atypia- to-carcinoma progression and is suspected to participate in the carcinogenesis and progression of invasive breast cancer, identifying this gene as an important biomarker for invasive breast cancers [102]. SLC3A2 was shown to play a role in aggressive Breast cancer subtypes driven by c-MYC and has been proposed as a potential poor prognostic marker for highly proliferative breast cancer subtypes [103]. P4HA1 expression has recently been shown to induce HIF-1α expression in TNBCs, revealing a novel HIF-1 regulation mechanism in TNBCs, and P4HA1 was also shown to promote chemoresistance by modulating HIF-1-dependent cancer cell stemness [104, 105]. SEPT7 along with SEPT2, cytoskeletal proteins evolutionary conserved from yeast to mammals, when silenced in MB231 cells, demonstrated inhibitory effects in cellular proliferation, apoptosis, migration, and invasion, strongly suggesting that SEPT7 is involved in breast carcinogenesis and can serve as therapeutic targets for highly invasive breast cancers [106]. IKBIP, a gene not commonly reported in cancer data, was recently shown to be highly involved in EMT and was associated with more aggressive phenotypes in gliomas [107]. Finally, down-regulation of TXNIP, demonstrated in breast, bladder, and gastric cancer was recently shown to be a common feature in human tumor xenograft models in which intra-tumoral leukocytes are responsible for the decreased expression of TXNIP, suggesting that TXNIP expression levels can be used to monitor the functional state of tumor-infiltrating leukocytes [108]. Finally, it needs to be emphasized that several of our identified biomarkers in this study are overlapped with previously identified DEGs/DEPs in several endothelial cell lines with disruption of the CSC [109, 110], confirming the importance of the CmP network in the tumorigenesis of CAW-TNBCs. Together, these findings provide possible mechanisms by which all of our identified intrinsic and inducible biomarkers in CAW-TNBC cells, under mPRs-specific steroid actions, could be acting to perturb the numerous tumorigenesis signaling pathways observed in our multi-omics analysis.

### Further validation of the CmP signaling network in cancer

PRG-mediated signaling has been shown to play a critical role in the progression of ovarian, as well as breast cancer [111, 112], and its ability to regulate specific functions in nPR(-) breast cancers implicates mPRs in breast cancer progression and initiation [30, 113]. Accumulating evidence over the years demonstrating the role of mPRs in reproductive tumor biology has grown exponentially as overexpression of mPRs has been associated with the development and progression of cervical and ovarian cancer [114–116] as well as breast cancers [32, 34]. Furthermore, mPRs are widely expressed in all breast cancer cell lines and biopsies [30, 32–34, 113].

Our previous studies investigating the role of mPRs in the CmP signaling network in AAW-TNBC progression, also provide additional support for these findings in which we observed expression levels of mPRs were altered at both the transcriptional and translational level in AAW-TNBC cells under mPRs-specific steroid actions, and that mPRs were localized within the nucleus and could participate in nucleo-cytoplasmic shuttling [31]. Additionally, we discovered that distinct RNA/protein expression patterns of key factors in the CmP network, under mPRs-specific steroid actions, allowed us to categorize TNBC cells into two subgroups: CAW-TNBCs, termed Type 2A and AAW-TNBCs, termed Type 2B [31].

In this study, we observed differential gene/protein expression of inducible biomarker CAV1, which is known to co-localize at caveolae with mPR*α* and epidermal growth factor receptor (EGFR) and interestingly, PRG has been shown to induce repression of EMT via caveola bound signaling complexes [32, 116]. Furthermore, mPR*α* has been shown to function as an essential mediator for PRG-induced inhibition on cell invasion and migration in basal phenotype breast cancer cells [117]. Together these results reinforce the overwhelming evidence of the involvement of the CmP signaling network in TNBC progression in both CAW and AAW [31].

## Conclusion

In our previous study, we defined the novel CmP signaling network in TNBC cells [31], which overlapped with our earlier defined CmPn network in nPR(+) breast cancer cells [25], and observed that mPRs-specific steroid actions have a positive effect on CSC protein expression in TNBCs with inducible patterns of expression in AAW-TNBC cells [31]. In this study, we were once again able to demonstrate alterations to key tumorigenesis pathways in CAW-TNBC cells, under mPRs-specific steroid actions, which were in parallel to our observations in AAW-TNBC cells under identical conditions [31]. These results suggest that even though TNBC diagnoses in AAW are associated with more aggressive forms of the disease [54–56], and experience a higher mortality rate [57], TNBC in CAW share the similar altered signaling pathways, under mPRs-specific steroid actions, demonstrating the aggressive nature of TNBCs regardless of racial differences. Utilizing expression and prognostic data, we were able to identify 21 clinically relevant intrinsic and inducible biomarkers for CAW-TNBCs (Table 1). This new set of potential prognostic biomarkers may revolutionize molecular mechanisms and currently known concepts of tumorigenesis in CAW-TNBCs, leading to hopeful new therapeutic strategies.

## Materials and Methods

### Cell culture and treatments

#### Cell culture and treatments

Cell culture and treatments were performed as previously described [31]. Briefly, nPR(-) MDA-MB-231 cells were cultured in RPMI1640 medium following manufacturer’s instructions (ATCC), and once cells reached 80% confluency, cells were starved with FBS-free RPMI1640 medium for 3 hrs, followed with either vehicle control (ethanol/DMSO, VEH) or combined steroids (MIF+PRG, 20 µM each). After cells were treated, they were harvested for RNA and protein extraction as described before [118, 119].

### RNA extraction, RT-qPCR, and RNA-seq for TNBC-CAW cells

RNA extraction was performed as previously described [31]. Briefly, total RNAs from MDA-MB-231 cells were extracted using TRIZOL reagent (Invitrogen) following the manufacturer’s protocol. The cell monolayer was rinsed with ice-cold PBS then lysed directly in a culture dish with 1 ml of TRIZOL reagent by scraping with a cell scraper. The cell lysate was aspirated and dispensed several times using a pipette and vortexed thoroughly. The purity and integrity of each RNA sample were analyzed using a Bioanalyzer (Agilent). All RNA-seq data were produced using Illumina HiSeq 2000; clean reads for all samples were over 99.5%; 60-80% of reads were mapped to respective reference genomes.

#### RNA-seq processing of files to assemble interactomes for TNBC-CAW cells

RNA-seq was performed as previously described [31]. Briefly, RNA-seq files were processed through paired-end (PE) sequencing with 100 bp reads (2X100) in Illumina HiSeq2000. The data consisted of 4 FASTQ files, 2 PE FASTQ files for each of the two groups: MDA-MB-231_Veh, MDA-MB- 231_MP treated 48hrs with both cohorts having biological and technical duplicates. RNA data were cleaned and ensured to pass quality control before performing bioinformatics analysis. After extracting the FASTQ files, they were aligned to the Human Genome Build 38 (GRCh38) using HISAT2 and then converted to SAM using SAMtools 1.9 and SAMtools quick check to ensure the files were complete. The SAM files were then converted to BAM files, sorted and Cufflinks was used to assemble transcripts and cuffmerge to merge all the transcript files. Differential gene expression was performed with CuffDiff followed by identifying differentially expressed genes between cohorts using a Python script and annotated in Excel. Identified significant differentially expressed genes were then filtered using our MD-MBA-468_MP treated 48hrs RNA-seq data [31] to obtain significant differentially expressed genes unique for MDA-MB-231cells (TNBC-CAW). The remaining differentially expressed genes after filtering were inputted into the PANTHER [120] classification system (GeneONTOLOGY) as well as iDEP [121] (Integrated Differential Expression and Pathway Analysis) program to generate heatmaps and build signaling networks with altered GO biological processes, GO molecular functions, PANTHER protein classes, and altered KEGG pathways.

### Protein extraction and proteomics analysis for TNBC-CAW cells

#### Protein Extraction and quality assessment

protein extraction for MDA-MB-231 cells was performed as previously described [31]. Briefly, MDA-MB-231 cells were harvested and lysed using a digital sonifier cell disruptor (Branson model 450 with model 102C converter and double step microtip) in ice-cold lysis buffer (50 mM Tris-HCl (pH 7.5), 150 mM NaCl, 0.5% NP-40 (Sigma), 50 mM sodium fluoride (Sigma), 1 mM PMSF (Sigma), 1 mM dithiothreitol (Invitrogen) and 1 EDTA-free complete protease inhibitor tablet (Roche)). Protein concentration was assessed using Qubit assay (Invitrogen). For proteomics analysis, 20-25 µg of proteins were prepped using the filter-aided sample preparation (FASP) method (Expedeon, San Diego, CA). Samples were reduced with 10 mM DTT for 30 min at RT, centrifuged on a spin filter at 14,000× g for 15 min at RT, and washed twice with 8 M urea in 50 mM Tris-HCl buffer. Samples were then washed 3x with 50 mM ammonium bicarbonate then digested with trypsin (Sigma-Aldrich, St. Louis, MO) and peptides eluted using 0.1% formic acid.

#### Liquid Chromatography-Tandem Mass Spectrometry (LC-MS/MS)

LC-MS/MS for MDA-MB-231 cells was performed as previously described [31]. Briefly, the cell lysates were generated from the two cohorts, MDA-MB-231_Veh and MDA-MB-231_MP treated 48hrs. All cohorts consisted of biological triplicates. After FASP preparation, four microliters of each digested sample (100 ng/µl) were loaded onto a 25-cm custom-packed porous silica Aqua C18, 125Å, 5 µm (Phenomenex) column. The porous silica was packed into a New Objective PicoTip Emitter, PF360-100-15-N-5, 15 ±1 µm, pre-equilibrated with 95% solvent A (100% water, 0.1% formic acid) and 5% solvent B (90% acetonitrile, 10% water, 0.1% formic acid) before injection of the digested peptides. LC separation of the peptides was performed on an Ultimate 3000 Dionex RSLC-nano UHPLC (Thermo Fisher Scientific), equilibrated with 95% solvent A and 5% solvent B (equilibration solution). Samples were loaded onto the column for 10 mins (flow rate of 0.5µL/min), before beginning the elution gradient. Solvent B was increased from 5%-35% over 85 min, followed by a sharp increase to 95% solvent B over 5 min. The plateau was maintained at 95% solvent B for 9 min, followed by a sharp decrease to 5% solvent B over 1 min. Peptides were analyzed using a Q Exactive Plus Hybrid Quadrupole-Orbitrap Mass Spectrometer (Thermo Fisher Scientific), equipped with a Nanospray Flex Ion Source (Thermo Fischer Scientific). Parameters for the mass spectrometer were as follows: full MS; the resolution was set at 70,000 and 17,500, for MS1 and MS2, respectively; AGC target was set at 3e6 and 1e5 for MS1 and MS2, respectively; Max IT at 50 ms; scan range from 350 to 1600 m/z dd-MS2; Max IT at 100 ms; isolation window at 3.0.

#### Proteomics processing of files for TNBC cells

Proteomic data analysis for MDA-MB-231 cells was performed as previously described [31]. Briefly, analysis was performed with Proteome Discoverer (PD) 2.1.1.21 (Thermo Fisher Scientific), using an FDR of 1%. The Human Database was downloaded in FASTA format on July 1, 2021, from UniProtKB; http://www.uniprot.org/. Common contaminants (trypsin autolysis fragments, human keratins, and protein lab standards) were included in the contaminants database [122]. The following parameters were used in the PD: HCD MS/MS; fully tryptic peptides only; up to 2 missed cleavages; parent-ion mass tolerance of 10 ppm (monoisotopic); fragment mass tolerance of 0.6 Da (in Sequest) and 0.02 Da (in PD 2.1.1.21) (monoisotopic). A filter of two high confidence peptides per protein was applied for identifications. PD dataset was further processed through Scaffold Q+ 4.8.2 (Proteome Software, Portland, OR). A protein threshold of 99%, peptide threshold of 95%, and a minimum number of 2 peptides were used for protein validation. Data were then exported to perSPECtives 3.0.0 (Proteome Software, Portland, OR) and used to validate and statistically compare protein identifications derived from MS/MS search results. Proteomic biological and technical samples were analyzed via Students *t-test* with a multiple test correction using Benjamini-Hochberg procedure and an FDR level of 0.05. Normalized weighted spectral counts were used for the comparisons. A cutoff of p≤0.05 was utilized to determine significance in vehicle and PRG+MIF treated comparisons.

#### Processing of proteomic files to assemble interactomes for TNBC cells

Bioinformatics data analysis for MDA-MB-231 cells was performed as previously described [31]. Briefly, a Python script was created to identify differentially expressed proteins in vehicle and PRG+MIF treated comparisons. These were annotated in Excel and identified significant differentially expressed proteins were then filtered using our MD-MBA-468_MP treated 48hrs proteomics data [31] to obtain significant differentially expressed proteins unique for MDA-MB-231cells (TNBC-CAW). The remaining differentially expressed proteins after filtering were inputted into the PANTHER [120] classification system (GeneONTOLOGY) as well as iDEP [121] (Integrated Differential Expression and Pathway Analysis) program to generate heatmaps and build signaling networks with altered GO biological processes, GO molecular functions, PANTHER protein classes, and altered KEGG pathways.

#### Omics analysis to assemble interactomes for TNBC-CAW cells at both the transcriptional and translational levels

A Python script was created to identify shared differentially expressed genes/proteins identified in MDA-MB-231_MP treated 48hrs compared to MDA-MB-231_Veh treated samples. These were annotated in Excel, and simple set operations were used to find the overlaps. The overlaps were then filtered using our MD-MBA-468_MP treated 48hrs proteomics/RNA-seq data [31] to obtain significant differentially expressed genes/proteins unique for MDA-MB-231cells (TNBC-CAW). The leftover genes/proteins were inputted into the PANTHER [120] classification system (GeneONTOLOGY) as well as iDEP [121] (Integrated Differential Expression and Pathway Analysis) program to generate heatmaps and build signaling networks with altered GO biological processes, GO molecular functions, PANTHER protein classes, and altered KEGG pathways. The goal was to identify similarities between MDA-MB-231_MP treated 48hrs samples (compared to vehicle samples) at both the transcriptional and translational levels. The agreements in differential expression were noted through the use of the Python comparison script.

### Prognostic effects for identified candidate biomarkers associated with a perturbed CmP signaling network in TNBC-CAW cells

#### Differential expression of candidate biomarkers using Microarray expression data

Expression analysis was performed as previously described [31]. Briefly, microarray data containing breast cancer tumors, with Progesterone Receptor (PR) status determined by IHC, were analyzed using kmplot software [49]. This resulted in 925 nPR(-)and 926 nPR(+) breast cancer samples that were used to obtain expression data for the candidate biomarkers identified in this study.

#### Differential expression of candidate biomarkers using RNA-seq data for AAW-TNBCs and CAW-TNBCs

Expression analysis for AAW-TNBCs and CAW-TNBCs was performed as previously described [31]. Briefly, to evaluate the basal expression of our identified candidate biomarkers in AAW and CAW TNBCs, we utilized publicly available RNA-seq data comparing expression levels in AAW-TNBCs and CAW-TNBCs [50]. This cohort contained 23 TNBC-AAW samples and 19 TNB-CAW samples that were used for the analysis of the candidate biomarkers identified in this study.

#### Construction of Kaplan-Meier (KM) survival curves for identified TNBC-CAW biomarkers

Construction of KM survival curves for CAW-TNBCs was performed as previously described [31]. Briefly, publicly available microarray data (22,277 probes) from 1,809 breast cancer patients were analyzed using kmplot software [49] integrating gene expression and clinical data simultaneously. Breast cancer patients were filtered to only analyze patient samples classified as ER(-)/nPR(-)/HER2(-)/TNBC subtype, which reduced our initial 1,809 patients to either 176-392 patients’ samples (depending on the biomarker analyzed). Logrank p-values, as well as hazard ratio (and 95% confidence intervals), were calculated by the software [49].

### Statistical Analysis

<u>For transcriptomics analysis</u>, all pairwise multiple comparison procedures were analyzed using Tukey and Student’s *t-test*. <u>For proteomics analysis</u>, all pairwise multiple comparison procedures were analyzed using Tukey and Student’s unpaired *t-test*. <u>For microarray/RNA-seq analysis,</u> Statistical significance was performed with Student’s *t*-test. All graphs, plots, and charts were constructed and produced by SigmaPlot 12.0 (Systat Software, Inc.) and GraphPad Prism 8.

## Acknowledgments

We wish to thank Victoria Reid, Chantal Alvarado, Yanchun Qu, Shen Sheng, Ahmed Badr, Junli Zhang, Amna Siddiqui, Pallavi Dubey, Saafan Malik, and Edna Lopez at Texas Tech University Health Science Center El Paso (TTUHSCEP) for their technical help during the experiments.

## Author contributions

JZ: Conceptualization, Writing-Reviewing and Editing; JAF: Investigation, Methodology, Software, Data curation, Validation, Writing-Original draft preparation, Writing-Reviewing and Editing, MB: Software, Data curation, Validation, Writing-Reviewing and Editing, BG: Software, Data curation, Validation;

