## Supplementary material for "Disrupting the CmP signaling network unveils novel biomarkers for triple negative breast cancer in Caucasian American women": Suppl Materials: Supplemental Legends-ALL.pdf

*Running Title:* Progesterone-mediated signaling in triple negative breast cancer (TNBC) cells

**All correspondence:
Jun Zhang, Sc.D., Ph.D.
Department of Molecular and Translational Medicine (MTM)
Texas Tech University Health Science Center El Paso
5001 El Paso Drive, El Paso, El Paso, TX 79905


**Suppl. Table 1: RAW RNAseq FASTQ files from MB231-specific Differentially Expressed Genes (DEGs).** We generated our RNAseq data specifically for MB231 (TNBC-CAW) cells by removing shared similarly altered DEGs overlapped with TNBC-AAW (MB468) cells. Comparison of DEGs (Column A) with technical p-values ≤ 0.05 from 48 hrs time points of TNBC-CAW cells treated with mPR specific steroids were used. Gene ID (Column B), Gene length (Column C),TNBC-CAW Vehicle treated averaged expression values (Column D), TNBC-CAW PRG+MIF treated averaged expression values (Column E), log2 Fold Change values (Column F), FDR (Column G), Up/Down regulation key (Column H), p-values (columns I) and signaling pathways data (Columns J-O) are displayed. Each treatment was run with biological and technical duplicates.

**Suppl. Table 2: RAW MS/MS data from MB231-specific Differentially Expressed Proteins (DEPs).** We filtered our MS/MS data specifically for MB231 (TNBC-CAW) cells by removing shared similarly altered DEPs overlapped with TNBC-AAW (MB468) cells. Comparison of DEPs (Column A) with technical p-values ≤ 0.05 from 48 hrs time points of TNBC-CAW cells treated with mPR specific steroids were used. Molecular weight of the proteins (Column B), p-values (column c), Fold change (Column D), log2 Fold Change values (Column E), TNBC-CAW Vehicle treated replicate expression values (Columns F-H), TNBC-CAW PRG+MIF treated replicate expression values (Columns I-K), and Up/Down regulation key (Column L) are displayed. Each treatment was run with biological and technical triplicates.

**Suppl. Table 3: RAW overlapped RNAseq and MS/MS data from MB231-specific Differentially Expressed Genes/Proteins (DEGs/DEPs).** We filtered our combined data specifically for MB231 (TNBC-CAW) cells by first finding similarly altered DEGs/DEPs between both omic data sets for MB231 (TNBC-CAW) cells, and then removing any overlapped DEGs/DEPs also identified in similarly altered DEGs/DEPs between both omic data sets for

TNBC-AAW (MB468) cells. **Tab 1-RNAseq**, identified DEGs (Column A) , Gene ID (Column B), Gene length (Column C),TNBC-CAW Vehicle treated averaged expression values (Column D), TNBC-CAW PRG+MIF treated averaged expression values (Column E), log2 Fold Change values (Column F), FDR (Column G), Up/Down regulation key (Column H), and p-values (columns I) are displayed. RNAseq samples were run with biological and technical duplicates. **Tab 2-Proteomics**, identification of DEPs (Column A), Molecular weight of the proteins (Column B), p-values (column c), Fold change (Column D), log2 Fold Change values (Column E), TNBC-CAW Vehicle treated replicate expression values (Columns F-H), TNBC-CAW PRG+MIF treated replicate expression values (Columns I-K), and Up/Down regulation key (Column L) are displayed. Proteomic samples were run with biological and technical triplicates.

**Suppl. Fig. 1. Expression and prognostic effects for identified candidate biomarkers utilizing microarray data of Breast cancer patient samples.** Publicly available microarray data (22,277 probes) from 1,809 breast cancer patients were analyzed using kmplot software to integrate gene expression and clinical data simultaneously to generate the displayed Kaplan-Meier survival curves. Breast cancer patients were filtered to only analyze patient samples classified as ER(-)/nPR(-)/HER2(-)/TNBC subtype, which reduced our 1,809 patients to 176/392 patient samples (depending on the probe). Results demonstrated **A-C)** significantly worst prognostic effects with our observed expression in our omics studies under a disrupted CmP network (mPR specific steroids) in CAW-TNBCs for **A1)** SLC3A2, **A2)** IKBIP, **A3)** ARRB1, **B1)** CD59, **B2)** GLS**, B3)** LDLR, **C1)** MCM4, **C2)** PTGES, and **C3)** TXNIP.**D-F).** Alternatively, results also demonstrated significantly better prognostic effects with our observed expression in our omics studies under a disrupted CmP network (mPR specific steroids) in CAW-TNBCs for **D1)** AK4, **D2)** CAV1, **D3)** F3, **D4)** TACSTD2, **E1)** LDHA**, E2)** PLOD2, **E3)** APAF1, **E4)** CDK2, **F1)** IQGAP3, **F2)** DRG1 (NDRG1), **F3)** P4HA1, and **F4)** SEPT7. Logrank p-values are calculated and displayed as well as hazard ratio (and 95% confidence intervals). The red line demonstrates high gene expression, while the black line demonstrates low gene expression.

**Suppl. Fig. 2. Non-significant Kaplan-Meier survival curves for overlapped DEGs/DEPs identified through our omics studies utilizing microarray data of TNBC patient samples.** Publicly available microarray data (22,277 probes) from 1,809 breast cancer patients was analyzed using kmplot software to integrate gene expression and clinical data simultaneously to generate the displayed Kaplan-Meier survival curves. Breast cancer patients were filtered to only analyze patient samples classified as ER(-)/PR(-)/HER2(-)/TNBC subtype, which reduced our 1,809 patients to 176/392 patient samples (depending on the probe). Results displayed demonstrate genes that did not quite reach significance ($p \leq 0.05$), eliminating them as identified candidate biomarkers. Logrank P values are calculated and displayed as well as hazard ratio (and 95% confidence intervals). Red line demonstrates high gene expression, while black line demonstrates low gene expression.

**Suppl. Fig. 3. Expression for identified candidate biomarkers utilizing microarray data of Breast cancer patient samples.** Publicly available microarray data (22,277 probes) from 1,809 breast cancer patients were analyzed using kmplot software and data was divided into two groups based on classic Progesterone Receptor (nPR) status determined by Immunohistochemistry (IHC); after filtering, there were 436-925 nPR(-) (depending on the probe) and 511-926 nPR(+) (depending on the probe) breast cancer samples. **(A-C)** Significant up-regulation of candidate biomarkers, in nPR(-) breast cancer tissue samples, compared to nPR(+) tissues, included **Ai)** AK4, **Aii)** APAF1, **Aiii)** CD59, **Aiv)** CDK2, **Bi)** LDHA, **Bii)** GLS**, Biii)** IQGAP3, **Biv)** PLOD2, **Ci)** NDRG1, **Cii)** SLC3A2, **Ciii)** P4HA1, and **Civ)** TACSTD2. **(D)** Significant down-regulation of candidate biomarkers in nPR(-) breast cancer tissue samples, compared to nPR(+) tissues, included **Di)** ARRB1, **Dii)** F3 **(Diii)** PTGES, and **(Div)** TXNIP. **(E)**

TNBC-CAW candidate biomarker expression was assessed using publicly available RNA-seq data for 23 TNBC-AAW samples and 19 TNBC-CAW samples to obtain expression data for TNBC-CAW candidate biomarkers in this study. Significantly decreased expression of candidate biomarkers in CAW-TNBCs, compared to AAW-TNBCs, included **Ei)** ARRB1, **Eii)** CDK2, **Eiii)** IQGAP3, **Eiv)** MCM4, **(Ev)** NDRG1, and **(Evi)** TACSTD2. Significantly increased expression of candidate biomarkers in CAW-TNBCs, compared to AAW-TNBCs, included **Fi)** CAV1, **Fii)** F3, **Fiii)** CD59, **Fiv)** LDHA, **(Fv)** GLS, **(Fvi)** PLOD2, **Fvii)** LDLR, **Fviii)** PTGES, **Fix)** IKBIP, and **(Fx)** TXNIP. Statistical significance was performed with student's *t*-test; *, **, *** above bars indicate P ≤ 0.05, 0.01, or 0.001, respectively.

**Suppl. Fig.4. Equal basal expression of TNBC-CAW candidate biomarkers, displaying significant survival curves, for breast cancer patient samples.** Microarray data containing breast cancer tumors, analyzed using kmplotsoftware, were divided into two groups based on Progesterone Receptor (PR) status determined by Immunohistochemistry (IHC); after filtering, there were 436-925 PR(-) (depending on biomarker) and 511-926 PR(+) (depending on biomarker) breast cancer samples. All panels illustrate genes that demonstrated equal expression in PR(-) breast cancer tissue samples, compared to PR(+) tissues. Additionally, all panels reflect genes that displayed significant differences in Kaplan-Meier survival curves. Statistical significance was performed with students t-test.

**Suppl. Fig.5. TNBC-CAW candidate biomarkers, displaying non-significant differential expression using RNA-seq data for AAW-TNBCs and CAW-TNBCs.** Publicly available RNAseq data for 23 TNBC-AAW samples and 19 TNBC-CAW samples were used to obtain expression data for TNBC-CAW candidate biomarkers in this study. All panels illustrate genes that demonstrated visual differences in expression between CAW and AAW TNBC-patients, but did not quite reach significance (p≤0.05). Additionally, all panels reflect genes that displayed significant differences in Kaplan-Meier survival curves. Statistical significance was performed with students t-test.

**Suppl. Fig.6. Shared Overlapped Differentially Expressed Genes/Proteins (DEGs/DEPs) among TNBC cells utilizing high-throughput RNA sequencing (RNAseq) and Proteomics.** In an effort to identify DEGs/DEPs shared between AAW-TNBCs and CAW-TNBCs we overlapped omics data between the two datasets (without filtering) to identify shared synchronous DEGs/DEPs to enhance our analysis. **A)** PCNA was analyzed (SRSF2 data not available) by a set of publicly available microarray data (22,277 probes) from 1,809 breast cancer samples using kmplot software to integrate gene expression and clinical data simultaneously to generate the displayed Kaplan-Meier survival curve results which demonstrated significant increased survival with our observed down-regulation for PCNA only. **B)** PCNA was further assessed (SRSF2 data not available) with two groups of breast cancer tissue samples, based on classic progesterone Receptor (nPR) status determined by Immunohistochemistry (IHC), to evaluate clinical expression levels focusing on 436-925 nPR(-) (depending on the probe) and 511-926 nPR(+) (depending on probe) breast cancer samples which demonstrated no significant differences for either gene **C)** PCNA and SRSF2 expression were further assessed in AAW-TNBC (n=23) and CAW-TNBC (n=19) tumor tissues which demonstrated no significant changes in gene expression between AAW-TNBCs and CAW-TNBCs (only PCNA results illustrated). **D)** Systems biology data for PCNA and SRSF2 is summarized, demonstrating PCNA as a potential intrinsic biomarker for general TNBCs, while SRSF2 appears to undergo potential feedback auto-regulation in both AAW-TNBC and CAW-TNBC cells under mPRs-specific steroid actions.

Suppl. Fig. 1.

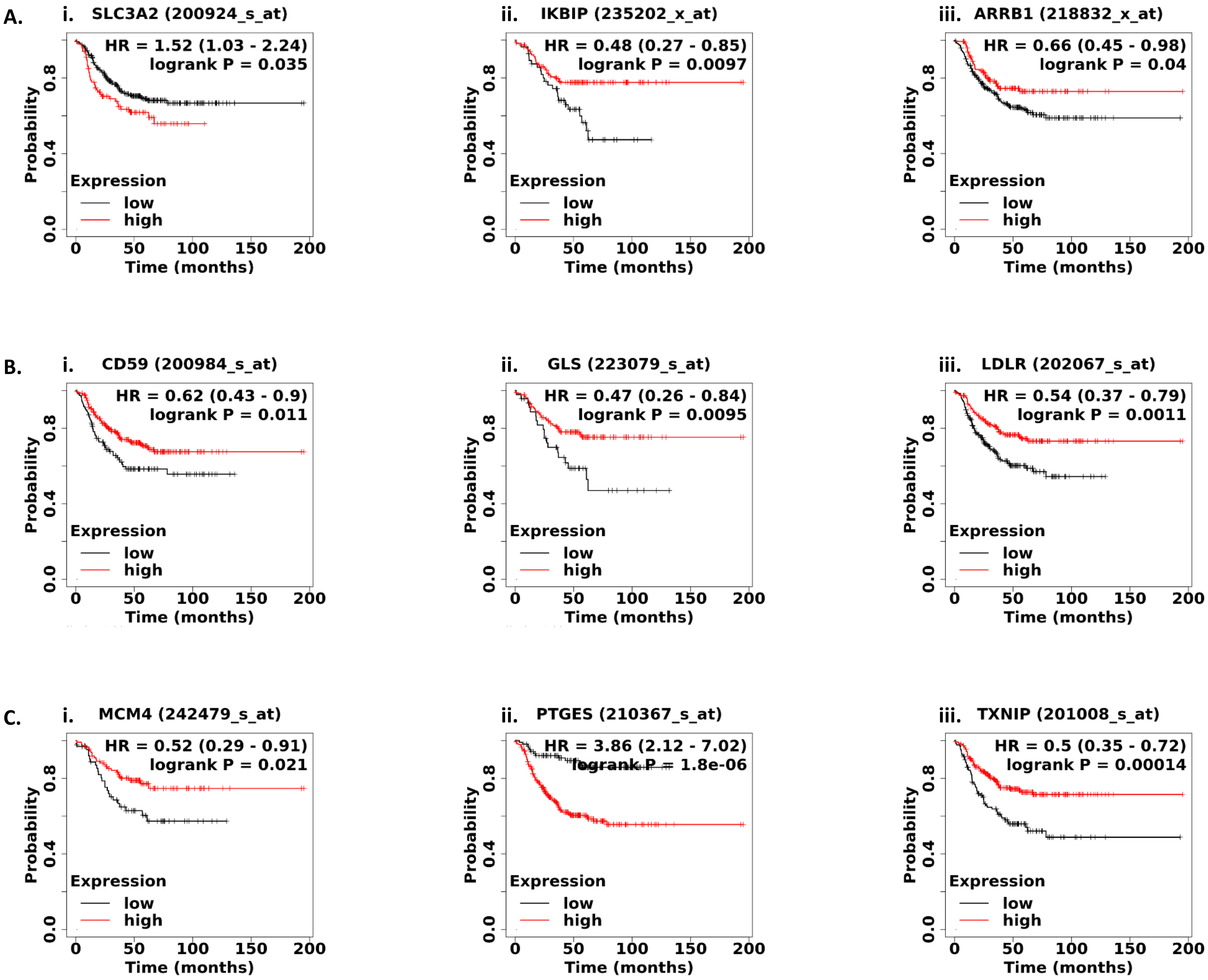
A.
i. SLC3A2 (200924_s_at)
HR = 1.52 (1.03 - 2.24)
logrank P = 0.035
ii. IKBIP (235202_x_at)
HR = 0.48 (0.27 - 0.85)
logrank P = 0.0097
iii. ARRB1 (218832_x_at)
HR = 0.66 (0.45 - 0.98)
logrank P = 0.04
B.
i. CD59 (200984_s_at)
HR = 0.62 (0.43 - 0.9)
logrank P = 0.011
ii. GLS (223079_s_at)
HR = 0.47 (0.26 - 0.84)
logrank P = 0.0095
iii. LDLR (202067_s_at)
HR = 0.54 (0.37 - 0.79)
logrank P = 0.0011
C.
i. MCM4 (242479_s_at)
HR = 0.52 (0.29 - 0.91)
logrank P = 0.021
ii. PTGES (210367_s_at)
HR = 3.86 (2.12 - 7.02)
logrank P = 1.8e-06
iii. TXNIP (201008_s_at)
HR = 0.5 (0.35 - 0.72)
logrank P = 0.00014
Probability
0.0
0.4
0.8
Expression
low
high
0
50
100
150
200
Time (months)

D.

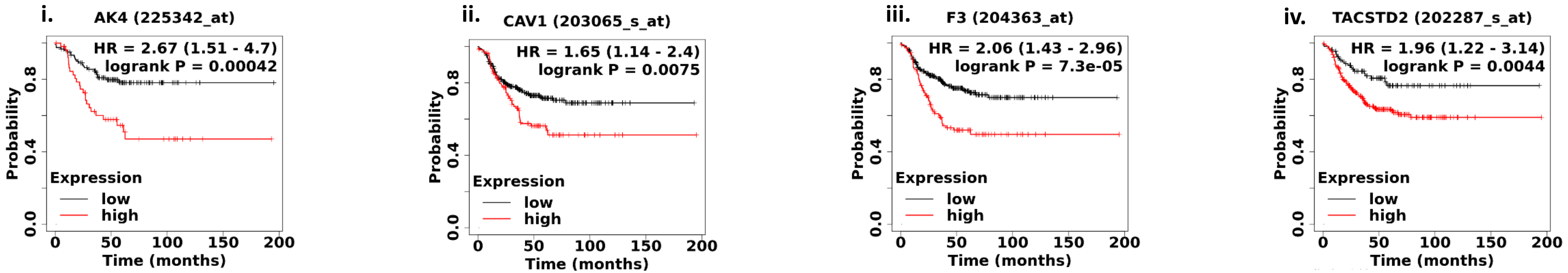
i. AK4 (225342_at)
HR = 2.67 (1.51 - 4.7)
logrank P = 0.00042
ii. CAV1 (203065_s_at)
HR = 1.65 (1.14 - 2.4)
logrank P = 0.0075
iii. F3 (204363_at)
HR = 2.06 (1.43 - 2.96)
logrank P = 7.3e-05
iv. TACSTD2 (202287_s_at)
HR = 1.96 (1.22 - 3.14)
logrank P = 0.0044
Probability
Time (months)
Expression
low
high

E.

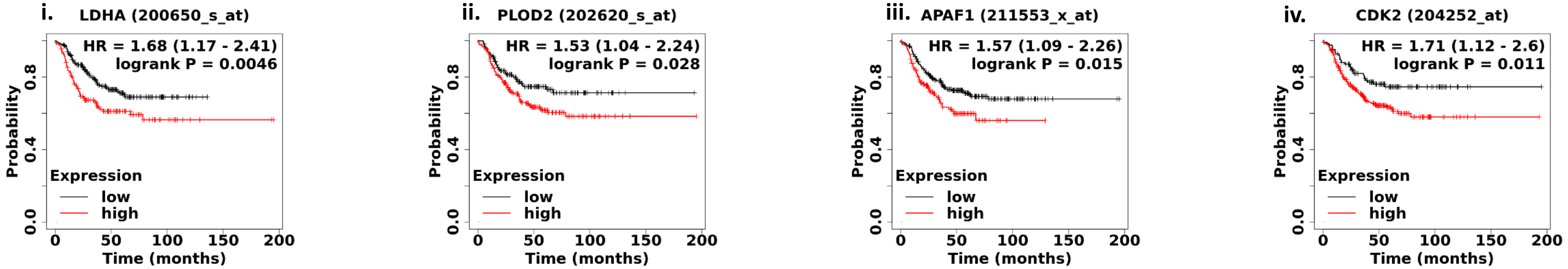
i. LDHA (200650_s_at)
HR = 1.68 (1.17 - 2.41)
logrank P = 0.0046
ii. PLOD2 (202620_s_at)
HR = 1.53 (1.04 - 2.24)
logrank P = 0.028
iii. APAF1 (211553_x_at)
HR = 1.57 (1.09 - 2.26)
logrank P = 0.015
iv. CDK2 (204252_at)
HR = 1.71 (1.12 - 2.6)
logrank P = 0.011
Probability
Time (months)
Expression
low
high

F.

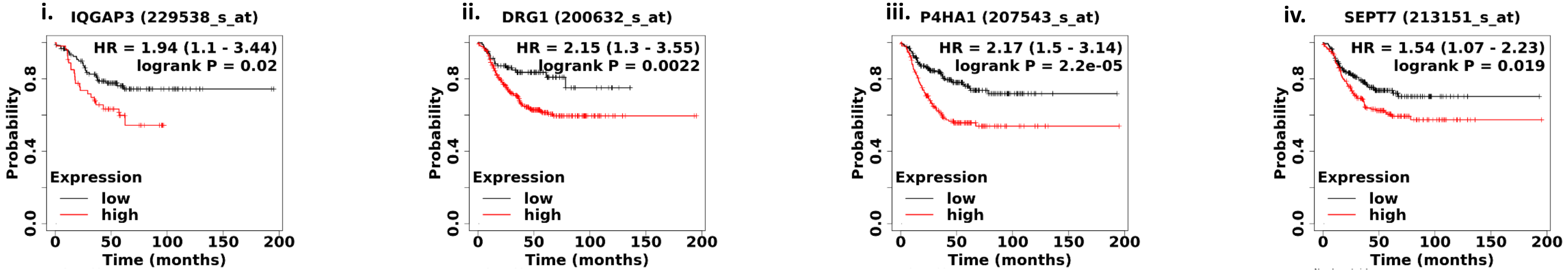
i. IQGAP3 (229538_s_at)
HR = 1.94 (1.1 - 3.44)
logrank P = 0.02
ii. DRG1 (200632_s_at)
HR = 2.15 (1.3 - 3.55)
logrank P = 0.0022
iii. P4HA1 (207543_s_at)
HR = 2.17 (1.5 - 3.14)
logrank P = 2.2e-05
iv. SEPT7 (213151_s_at)
HR = 1.54 (1.07 - 2.23)
logrank P = 0.019
Probability
Time (months)
Expression
low
high

### Suppl. Fig. 2

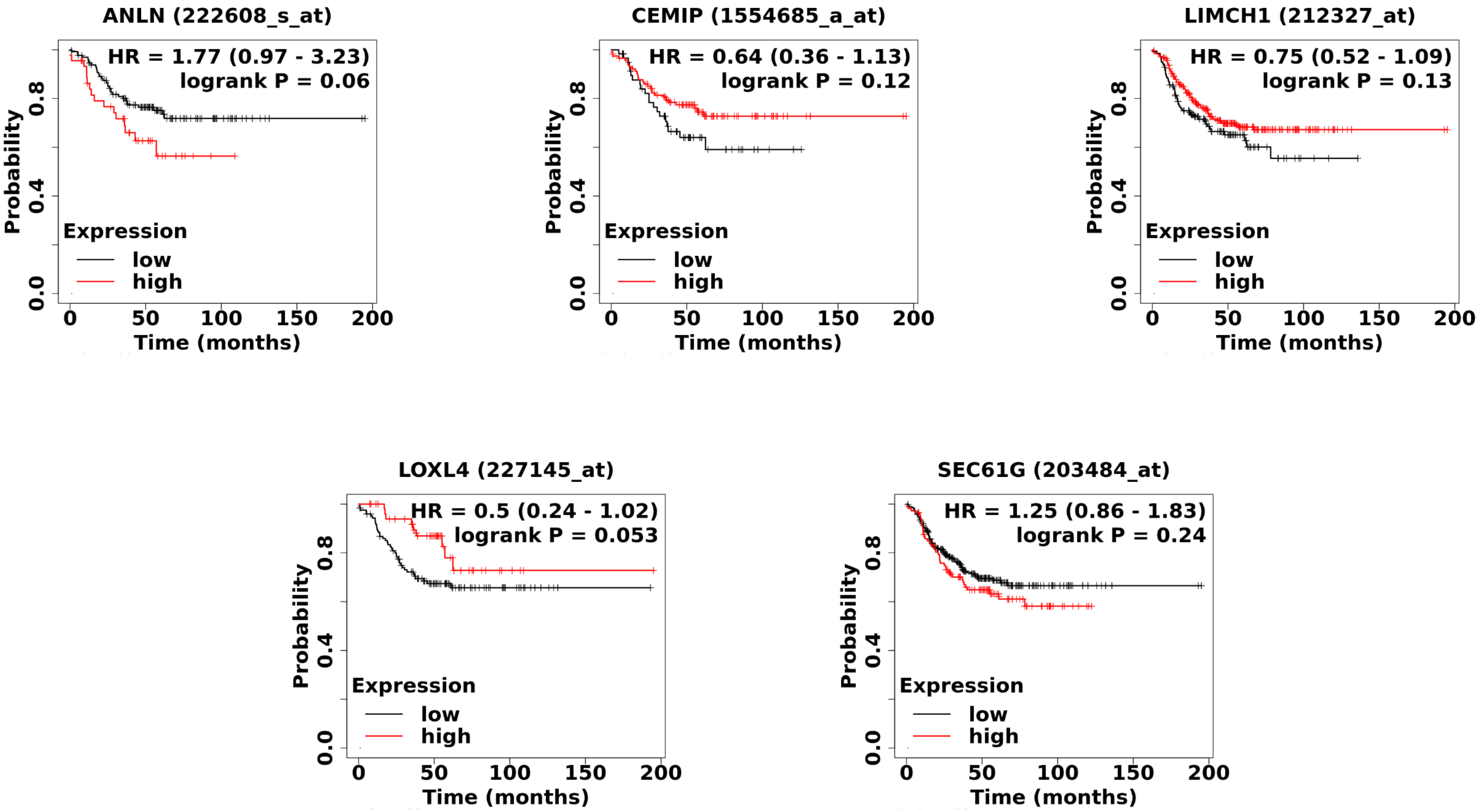
ANLN (222608_s_at)
HR = 1.77 (0.97 - 3.23)
logrank P = 0.06
CEMIP (1554685_a_at)
HR = 0.64 (0.36 - 1.13)
logrank P = 0.12
LIMCH1 (212327_at)
HR = 0.75 (0.52 - 1.09)
logrank P = 0.13
LOXL4 (227145_at)
HR = 0.5 (0.24 - 1.02)
logrank P = 0.053
SEC61G (203484_at)
HR = 1.25 (0.86 - 1.83)
logrank P = 0.24
Probability
0.0
0.4
0.8
Expression
low
high
0
50
100
150
200
Time (months)

Suppl. Figure 3.

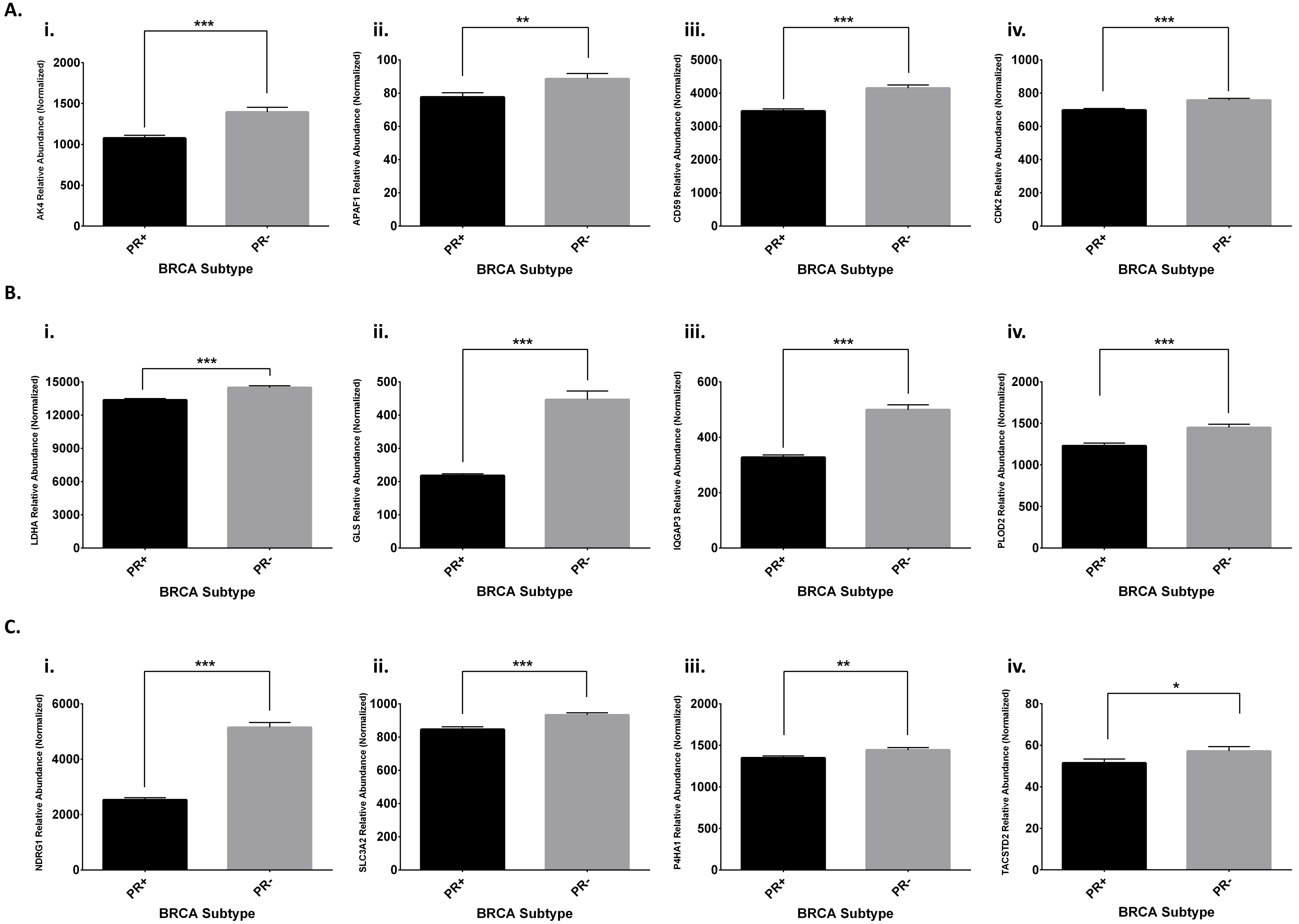

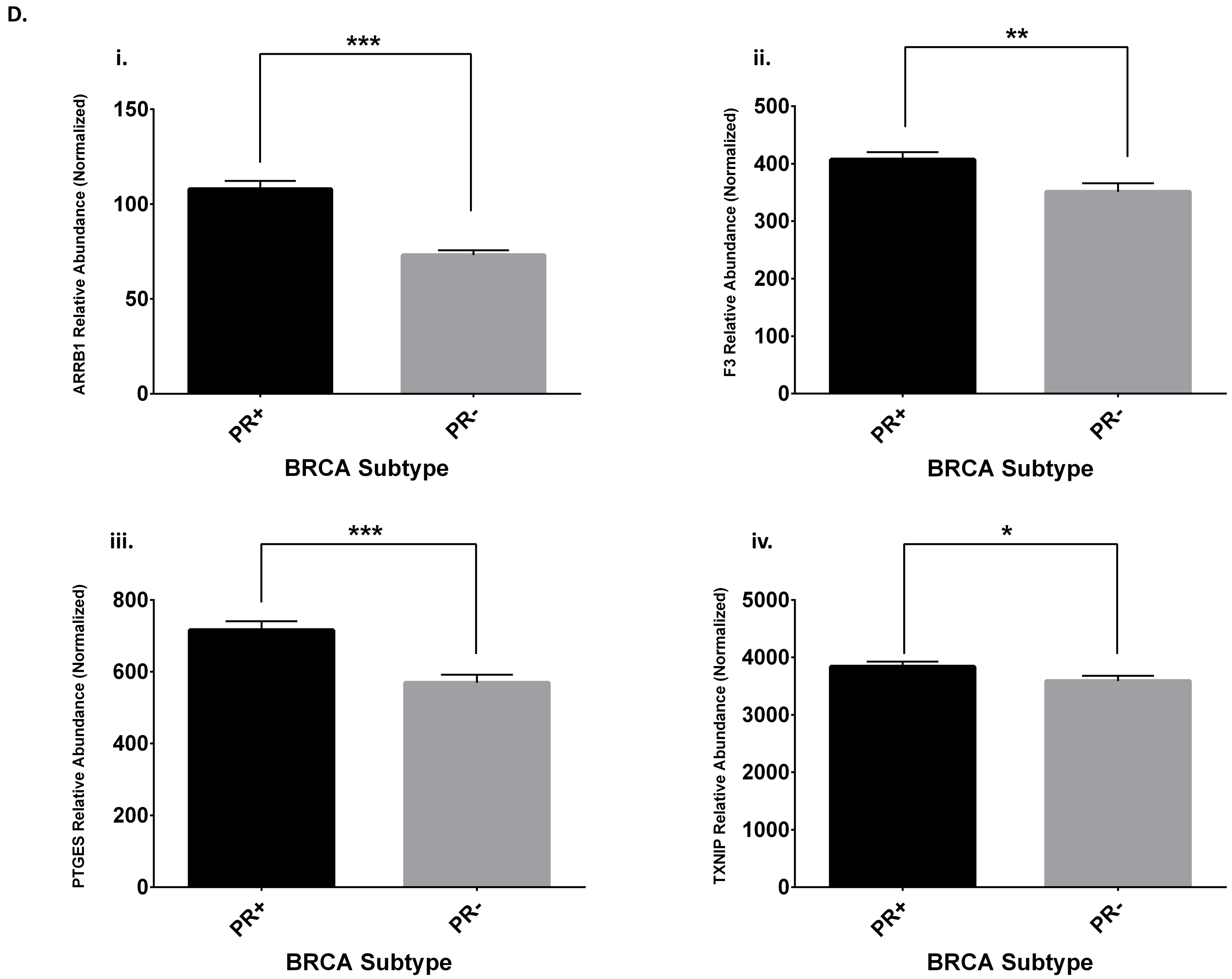
D.
i.
***
ARRB1 Relative Abundance (Normalized)
150
100
50
0
PR+
PR-
BRCA Subtype
ii.
**
F3 Relative Abundance (Normalized)
500
400
300
200
100
0
PR+
PR-
BRCA Subtype
iii.
***
PTGES Relative Abundance (Normalized)
800
600
400
200
0
PR+
PR-
BRCA Subtype
iv.
*
TXNIP Relative Abundance (Normalized)
5000
4000
3000
2000
1000
0
PR+
PR-
BRCA Subtype

E.

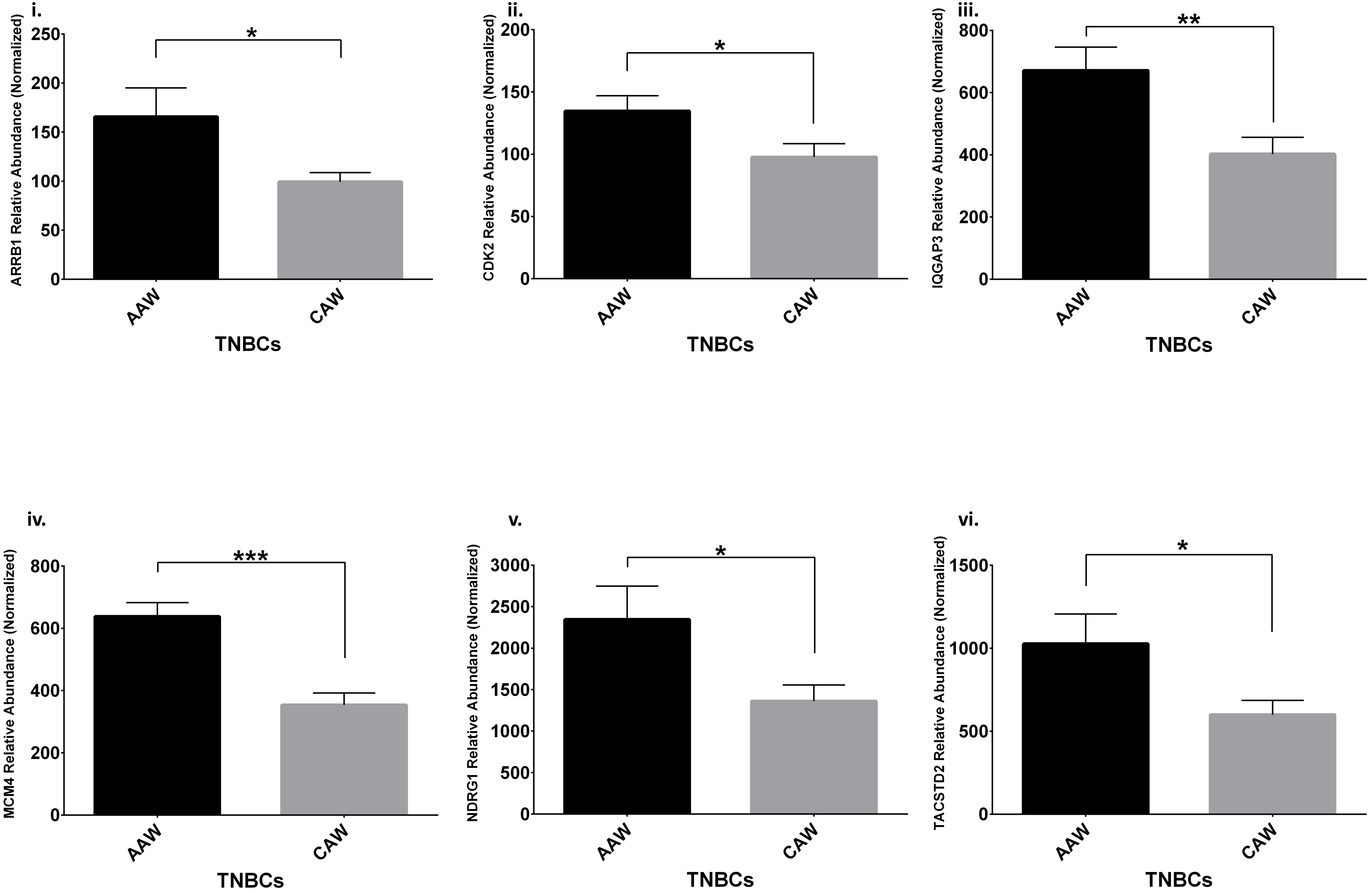

i.
ARRB1 Relative Abundance (Normalized)
250
200
150
100
50
0
*
AAW
CAW
TNBCs
ii.
CDK2 Relative Abundance (Normalized)
200
150
100
50
0
*
AAW
CAW
TNBCs
iii.
IQGAP3 Relative Abundance (Normalized)
800
600
400
200
0
**
AAW
CAW
TNBCs
iv.
MCM4 Relative Abundance (Normalized)
800
600
400
200
0
***
AAW
CAW
TNBCs
v.
NDRG1 Relative Abundance (Normalized)
3000
2500
2000
1500
1000
500
0
*
AAW
CAW
TNBCs
vi.
TACSTD2 Relative Abundance (Normalized)
1500
1000
500
0
*
AAW
CAW
TNBCs

F.

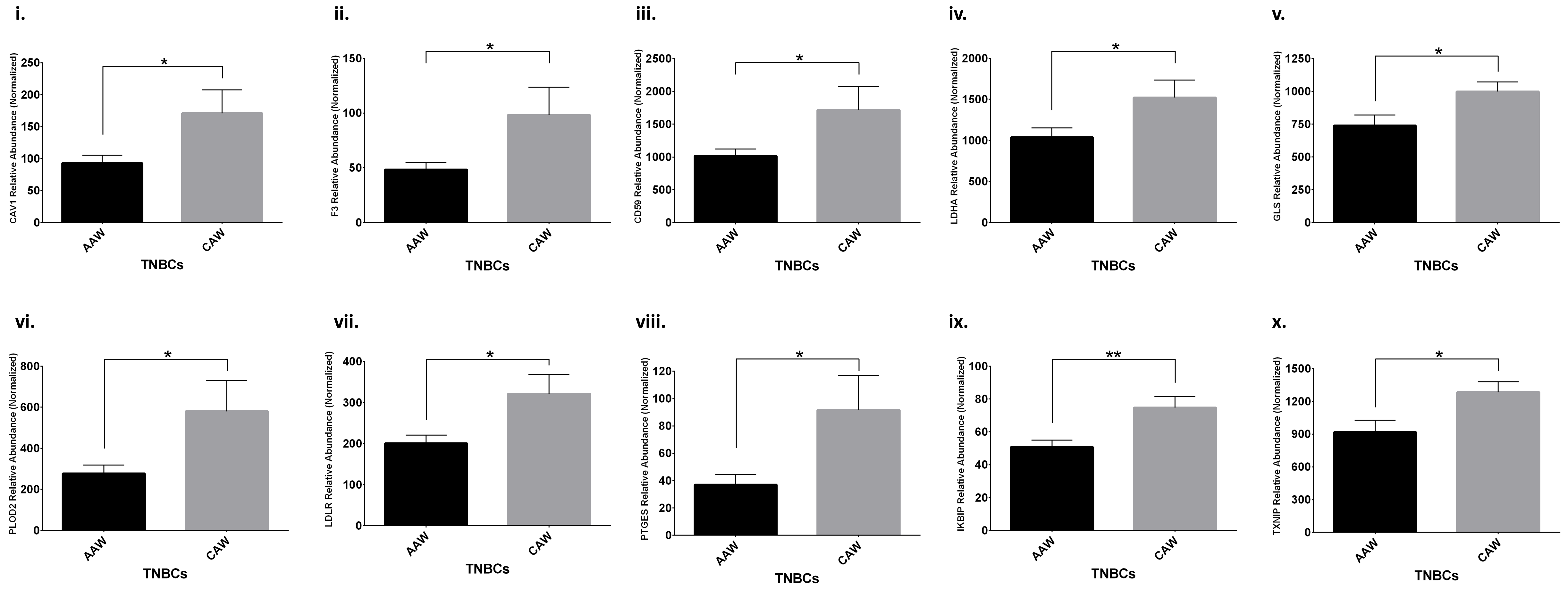

i.
CAV1 Relative Abundance (Normalized)
0
50
100
150
200
250
*
AAW
CAW
TNBCs
ii.
F3 Relative Abundance (Normalized)
0
50
100
150
*
AAW
CAW
TNBCs
iii.
CD59 Relative Abundance (Normalized)
0
500
1000
1500
2000
2500
*
AAW
CAW
TNBCs
iv.
LDHA Relative Abundance (Normalized)
0
500
1000
1500
2000
*
AAW
CAW
TNBCs
v.
GLS Relative Abundance (Normalized)
0
250
500
750
1000
1250
*
AAW
CAW
TNBCs
vi.
PLOD2 Relative Abundance (Normalized)
0
200
400
600
800
*
AAW
CAW
TNBCs
vii.
LDLR Relative Abundance (Normalized)
0
100
200
300
400
*
AAW
CAW
TNBCs
viii.
PTGES Relative Abundance (Normalized)
0
20
40
60
80
100
120
*
AAW
CAW
TNBCs
ix.
IKBIP Relative Abundance (Normalized)
0
20
40
60
80
100
**
AAW
CAW
TNBCs
x.
TXNIP Relative Abundance (Normalized)
0
300
600
900
1200
1500
*
AAW
CAW
TNBCs

Suppl. Fig. 4.

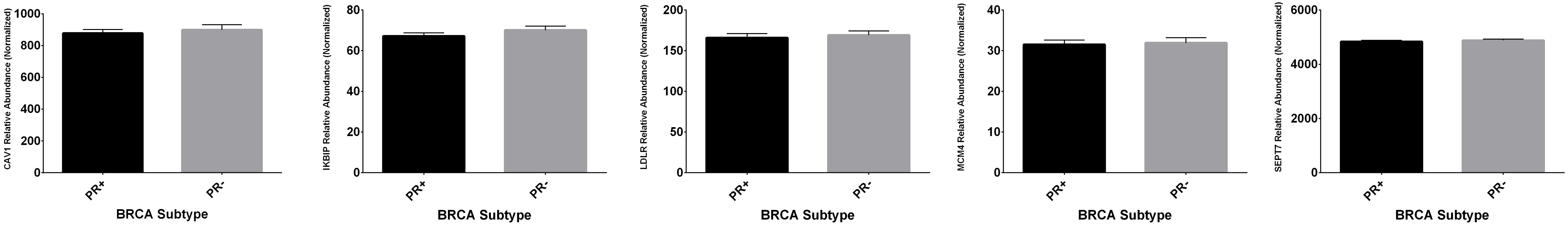
CAV1 Relative Abundance (Normalized)
1000
800
600
400
200
0
PR+
PR-
BRCA Subtype
IKBIP Relative Abundance (Normalized)
80
60
40
20
0
PR+
PR-
BRCA Subtype
LDLR Relative Abundance (Normalized)
200
150
100
50
0
PR+
PR-
BRCA Subtype
MCM4 Relative Abundance (Normalized)
40
30
20
10
0
PR+
PR-
BRCA Subtype
SEPT7 Relative Abundance (Normalized)
6000
4000
2000
0
PR+
PR-
BRCA Subtype

**Suppl. Fig. 5**

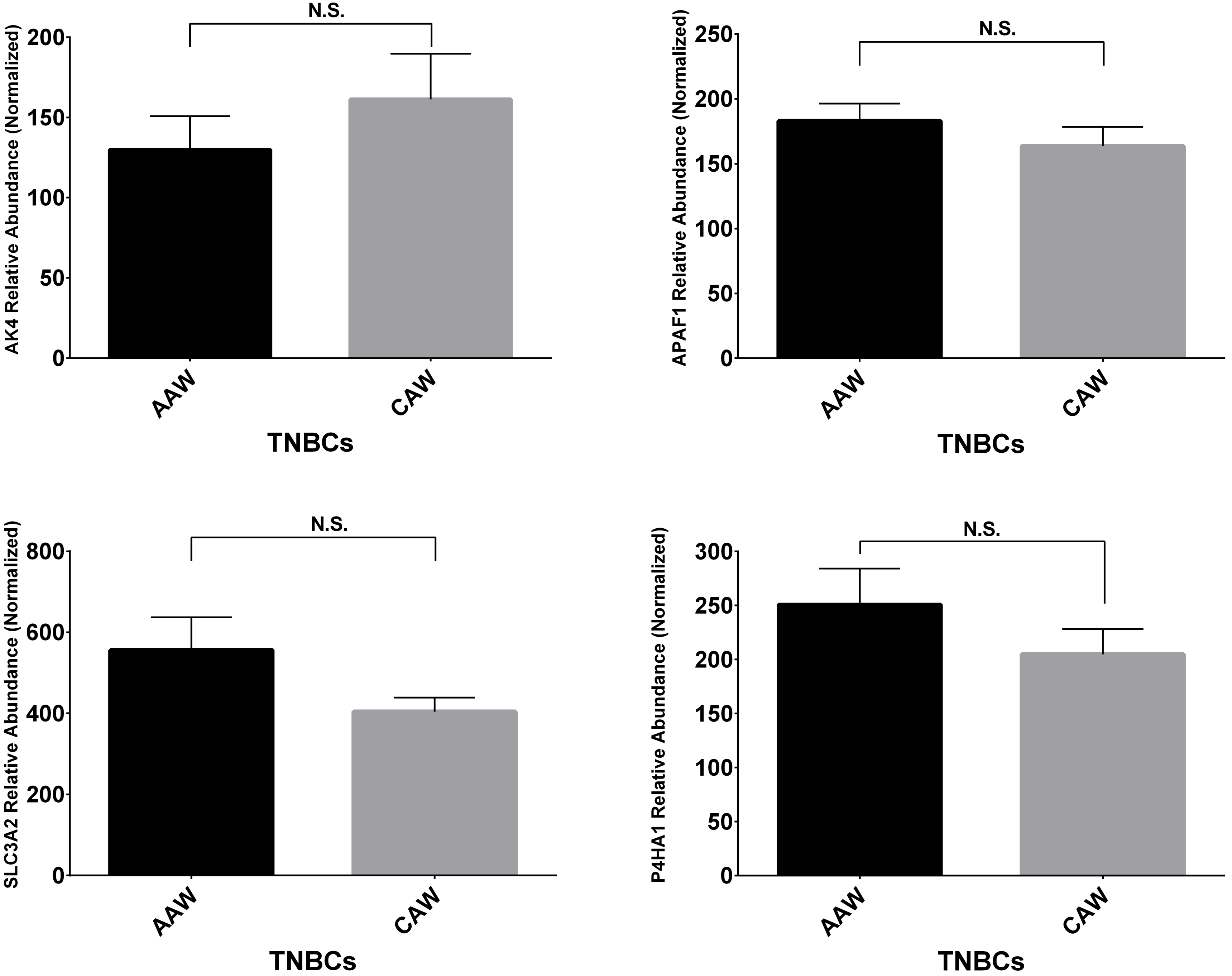
N.S.
AK4 Relative Abundance (Normalized)
200
150
100
50
0
AAW
CAW
TNBCs
N.S.
APAF1 Relative Abundance (Normalized)
250
200
150
100
50
0
AAW
CAW
TNBCs
N.S.
SLC3A2 Relative Abundance (Normalized)
800
600
400
200
0
AAW
CAW
TNBCs
N.S.
P4HA1 Relative Abundance (Normalized)
300
250
200
150
100
50
0
AAW
CAW
TNBCs

Suppl. Fig. 6

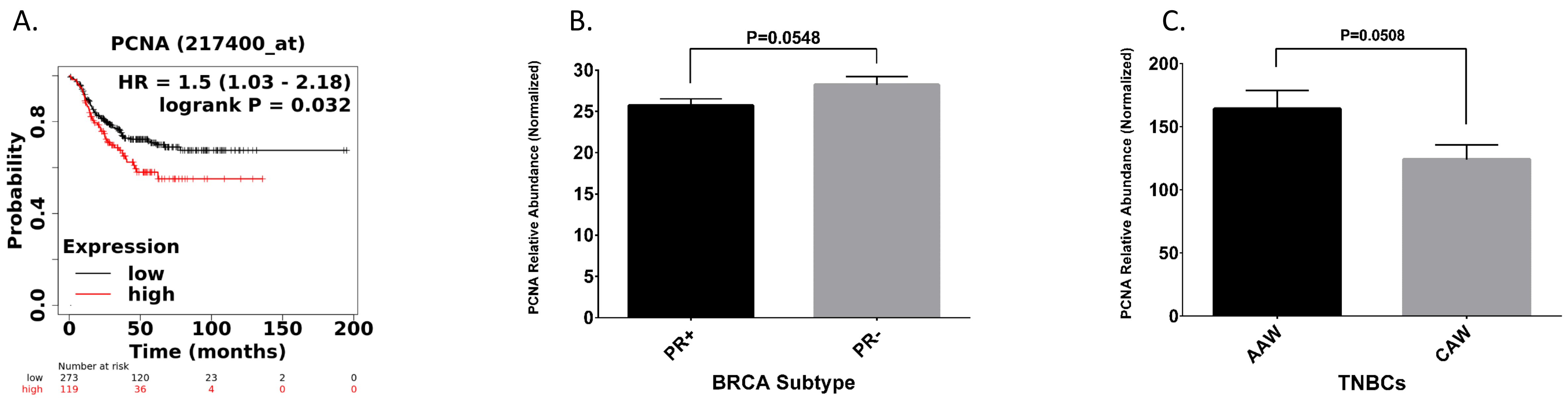


D.

| BIOMARKERS (ALTERNATE ID) | DEGs-TNBC AAW | DEPs-TNBC AAW | TNBC-AAW N=23 patients (RNA) | BIOMARKERS (ALTERNATE ID) | DEGs-TNBC CAW | DEPs-TNBC CAW | TNBC-CAW N=19 patients (RNA) | K-M Survival curve TNBC PATIENTS | Expression differences N= 925 PR(-) patients; N=926 PR(+) patients |
|---|---|---|---|---|---|---|---|---|---|
| PCNA | | | N.S. | PCNA | | | N.S. | I.S. N=392 | N.S. |
| SRSF2 | | | N.S. | SRSF2 | | | N.S. | N.A. | N.A. |
